# Spatiotemporal Control of Actomyosin Contractility by MRCKβ Signaling Drives Phagocytosis

**DOI:** 10.1101/2021.03.25.436833

**Authors:** Ceniz Zihni, Anastasios Georgiadis, Conor M. Ramsden, Elena Sanchez-Heras, Britta Nommiste, Olha Semenyuk, James W.B. Bainbridge, Peter J. Coffey, Alexander J. Smith, Robin R. Ali, Maria S. Balda, Karl Matter

**Affiliations:** Gene and Cell Therapy Group, UCL Institute of Ophthalmology, University College London; NIHR Biomedical Research Centre at Moorfields Eye Hospital NHS Foundation Trust and UCL Institute of Ophthalmology, City Road, London, EC1V 2PD, UK

**Keywords:** MRCK, Rho GTPases, Cdc42, actin dynamics, actomyosin contractility, membrane morphodynamics, phagocytosis, MerTK receptor, Fc-Receptor, membrane trafficking, adhesion, RPE, macrophages

## Abstract

Phagocytosis requires myosin-generated contractile force to regulate actin dynamics. However, little is known about the molecular mechanisms that guide this complex morphodynamic process. Here we show that particle binding to Mer Tyrosine Kinase (MerTK), a widely expressed phagocytic receptor, stimulates phosphorylation of the Cdc42 GEF Dbl3 in the retinal pigment epithelium (RPE), triggering activation of MRCKβ and its co-effector N-WASP that cooperate to deform the membrane into cups. Continued MRCKβ activity then drives recruitment of a mechanosensing bridge enabling transmission of the cytoskeletal force required for cup closure and particle internalization. MRCKβ is also required for Fc receptor-mediated phagocytosis by macrophages. *In vivo*, MRCKβ is essential for RPE phagocytosis of photoreceptor debris and, hence, retinal integrity. MerTK-independent activation of MRCKβ signaling by a phosphomimetic Dbl3 mutant rescues phagocytosis in retinitis pigmentosa RPE cells lacking functional MerTK. Thus, conserved MRCKβ signaling controls spatiotemporal regulation of actomyosin contractility to guide actomyosin dynamics-driven phagocytosis and represents the principle phagocytic effector pathway downstream of MerTK.

## INTRODUCTION

Phagocytosis is an ancient biological process that is essential for the functions of a variety of different cell types ranging from macrophages and microglia to specialized epithelia such as the retinal pigment epithelium (RPE) (Chaitin and Hall, 1983) (Inana et al., 2018) (Sparrow et al., 2010). Phagocytosis is triggered by cell surface receptors binding to a target particle, often in concert with co-receptors, stimulating sequential morphological changes leading to particle internalization (Freeman and Grinstein, 2014; Jaumouille et al., 2019; Swanson, 2008). Some phagocytes, such as macrophages, express a repertoire of specialized receptors with different specificities, such as Fc receptor (FcR), to phagocytose different types of particles. Other phagocytic receptors are widely expressed and are found in most phagocytic cells. This includes the TAM receptor MerTK that regulates phagocytosis by phagocytes of the immune system as well as phagocytic cells such as microglia, Schwann cells and the RPE. Hence, deregulation of MerTK-regulated phagocytosis has been linked to many diseases, ranging from cancer and rheumatoid arthritis to neurodegenerative conditions and blindness (Brosius Lutz et al., 2017) (Myers et al., 2019) (Haukedal and Freude, 2019) (D’Cruz et al., 2000). However, the downstream signaling mechanisms by which MerTK drives membrane morphodynamics to internalize particles is unknown.

In the eye, phagocytosis by the RPE is essential for clearing shed photoreceptor outer segment (POS) debris as part of a diurnal renewal process of photoreceptor membranes damaged by light (Young, 1967). Particle binding to the RPE membrane requires αvβ5 integrin (Finnemann et al., 1997; Finnemann and Rodriguez-Boulan, 1999; Lin and Clegg, 1998; Miceli et al., 1997) and MerTK, with the integrin being required for binding but not internalization of POS (Chaitin and Hall, 1983; D’Cruz et al., 2000; Edwards and Szamier, 1977; Mullen and LaVail, 1976; Ramsden et al., 2017). Internalization is stimulated by MerTK, a step essential for phagocytosis; hence, mutations inactivating MerTK lead to retinitis pigmentosa, an inherited form of retinal degeneration (Sparrow et al., 2010). Defects in phagocytosis also lead to age-related diseases in the eye due to deregulation of mechanisms that are poorly understood (Inana et al., 2018; Sparrow et al., 2010).

Despite the diversity of phagocytic receptors, cell types, and target particles’ properties, particle engulfment always requires a dynamic actin cytoskeleton. Engulfment is generally initiated by induction of membrane protrusions, such as the pseudopod-like structures induced by Cdc42 in FcR-mediated phagocytosis in macrophages, that then mature into a phagocytic cup wrapping the particle, which is followed by cup closure and internalization (Barger et al., 2020). The RPE membrane has been shown to generate pseudopod-like protrusions (Jiang et al., 2015), topologically similar to protrusions generated in FcR-mediated Cdc42-driven macrophage phagocytosis that are thought to require actomyosin contractility to remodel the membrane to wrap and phagocytose targets (Barger et al., 2020). However, the molecular mechanisms that spatially and temporally regulate actomyosin contractility and couple it to distinct morphological transformations to control particle internalization are not known in any phagocytic cell-type.

Here, we find that the myosin-II activator MRCKβ is required for spatiotemporal regulation of actomyosin contractility to drive phagocytosis and identify the engulfment pathway downstream of MerTK receptor activation. MerTK activates the Cdc42 guanine nucleotide exchange factor (GEF) Dbl3 through Src dependent phosphorylation. Locally activated Cdc42 then stimulates two effectors, N-WASP and MRCKβ. N-WASP stimulates actin-based pseudopod-like protrusions, and MRCKβ activates myosin-II to limit actin polymerization and control deformation of pseudopods into cups. MRCKβ-driven contractility also stimulates assembly of a mechanosensory bridge consisting of activated FAK, talin and viculin, and integrin clustering to guide membrane wrapping around particles. Finally, MRCKβ drives completion of cup closure and particle internalization, a function of MRCKβ also required for FcR-mediated macrophage phagocytosis. The functional importance of the discovered mechanism is further supported by its requirement to maintain retinal integrity *in vivo*. Direct activation by expression of a phosphomimetic Dbl3 mutant that circumvents the requirement for MerTK rescues phagocytosis in iPSC-derived retinitis pigmentosa RPE cells, indicating that the principal function of MerTK in phagocytosis is activation of the MRCKβ pathway.

## RESULTS

### Conserved cortical Cdc42 signaling drives phagocytosis in the RPE

Phagocytosis in the RPE involves the generation of pseudopod-like protrusions with a morphology similar to those in macrophages induced by particle binding to Fc-Receptors (Jiang et al., 2015). Macrophages require Cdc42 signaling to initiate phagocytosis (Barger et al., 2020). The apical membrane of epithelia is a domain enriched in evolutionarily conserved Cdc42 signaling components; however, their postmitotic functions once cells have polarized are not known. Therefore, we tested whether Cdc42, its apical GEF Dbl3 and downstream effectors including the myosin II activator MRCKβ are involved in phagocytosis (Kumfer et al., 2010; Marston et al., 2016; Zihni et al., 2016; Zihni et al., 2014; Zihni et al., 2017b) (Zihni and Terry, 2015) (Zihni et al., 2017a). Depletion of both Dbl3 and Cdc42 in confluent polarized porcine primary RPE cells resulted in inhibition of POS internalization (Fig.1a-e, Fig.S1a-c). POS binding to the RPE was not affected, indicating normal receptor expression (Fig.1b,e). Addition of POS to Dbl3 or Cdc42 depleted cells frequently resulted in apical F-actin-based membrane protrusions that failed to wrap around POS particles (Fig.S1a,b-right panels). Since we had used an incubation time sufficient for completion of the phagocytic process (1h POS binding followed by a 2h chase), this suggests that loss of Cdc42 is accompanied by a defect in POS-induced actin dynamics required for phagocytosis. POS binding stimulated local Cdc42 activation at attachment sites following short 20min incubations, which was strongly attenuated by knockdown of Dbl3, indicating that Dbl3 activates Cdc42 at the onset of POS/membrane binding (Fig.1f; Fig.S1d).

**Figure 1.**
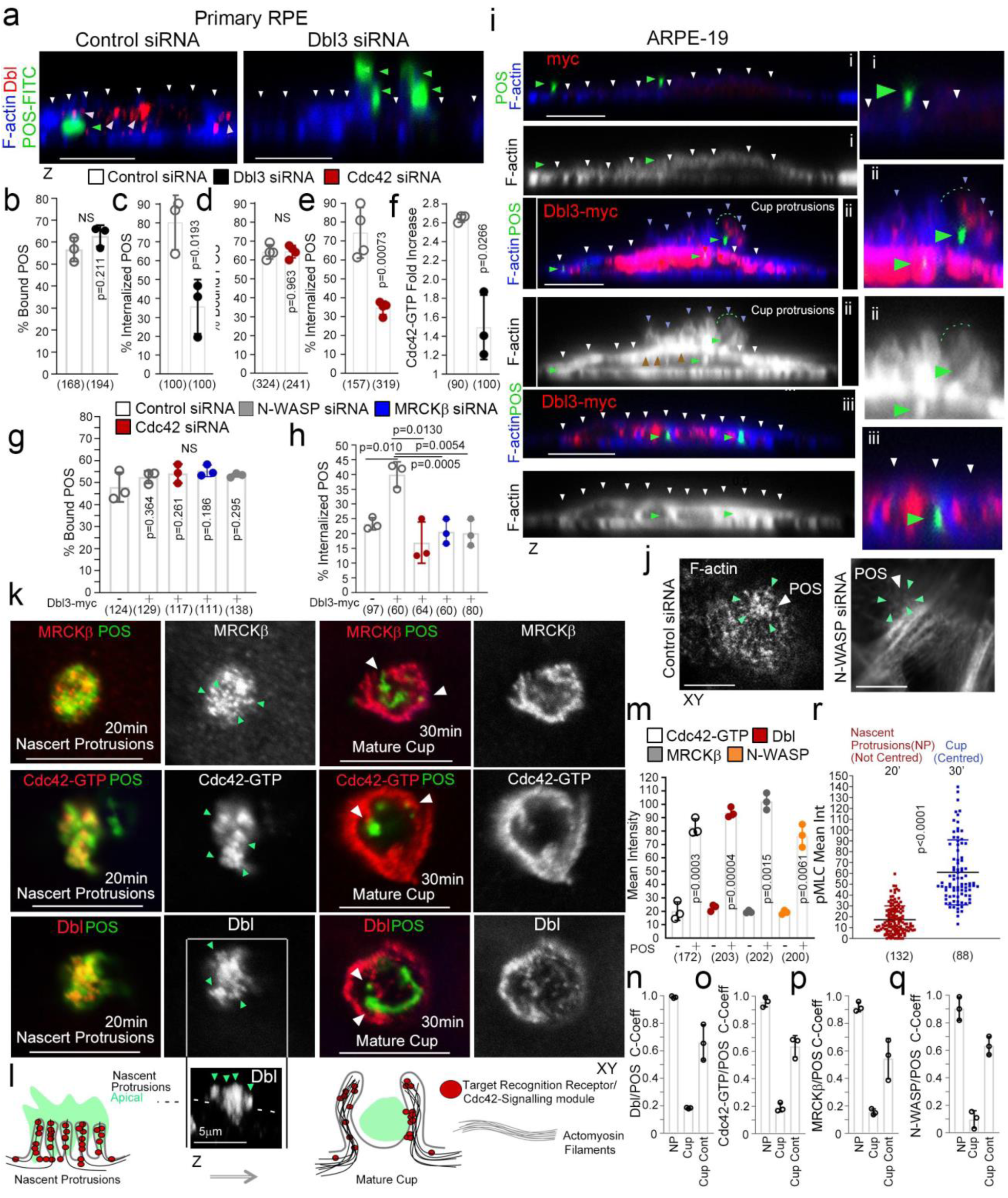
Apical Cdc42-signaling drives cup formation and engulfment in the RPE. **a**-**e**, siRNA-mediated knockdown of Dbl3 or Cdc42 in primary porcine RPE results in inhibition of phagocytosis. **f**, siRNA-mediated knokdown of Dbl3 in primary porcine RPE inhibits Cdc42 activation. **g-i**, Exogenous expression of Dbl3-myc in ARPE-19 cells stimulates increased POS-induced pseudopod and cup assembly, and phagocytosis via Cdc42, MRCKβ, and N-WASP signaling. White arrows highlight the apical F-actin cortex, green arrows position of POS and red arrows highlight Dbl3-myc localization at the tips of POS-stimulated cups. **j**, N-WASP siRNA-mediated knockdown in Dbl3-myc expressing ARPE-19 cells followed by 30min of POS stimulation inhibits F-actin assembly at the vicinity of POS adhesion. White arrows indicate position of POS and green arrows position of F-actin. **k-q**, POS-membrane contact rapidly induces membrane protrusions enriched in the Cdc42 GEF Dbl3, active Cdc42-GTP and the Cdc42 effectors MRCKβ and N-WASP. Note, nascent protrusions display almost complete co-localization of Cdc42-signaling components with POS, as determined by the Pearson’s Coefficient, but in cup conformation co-localization only occurs at discrete contact sites (highlighted by white arrows) within the cup-centered POS interface. The schematic illustration, **l**, outlines the Cdc42 signaling machinery that is recruited to nascent adhesions to induce nascent protrusions and subsequent membrane deformation into cups. In the cup conformation, the Cdc42 signaling machinery localizes at the POS-inner cup interface. **r**, Quantification of pMLC staining reveals progressively increasing myosin-II activation during nascent protrusion to cup maturation. Protein staining was quantified as mean intensity. Scale bars represent 10μm, unless highlighted otherwise. All quantifications are based on n=3 independent experiments, and show the data points, means ±1SD, the total number of cells analyzed for each type of sample across all experiments, and p-values derived from t-tests.

To examine how Cdc42 signaling initiates particle engulfment, we employed ARPE-19 cells, which normally phagocytose very slowly, as a gain of function model (Ablonczy et al., 2011). Exposure of ARPE-19 cells to POS-FITC for 1h followed by a 1h chase resulted in efficient binding of POS-FITC to the apical membrane, but protrusion induction and particle internalization were inefficient (Fig.1g-i; Fig.S1e). In contrast, exogenous expression of Dbl3-myc led to prominent POS-dependent nascent protrusions, phagocytic cups, and POS internalization (Fig.1h,i; Fig.S1e). Protrusions induced by Dbl3 were specifically activated by POS-membrane binding as they were not observed by expression of exogenous Dbl3 alone (Fig.S1e-right panels). Dbl3-myc localized along these nascent protrusions up to the tips (Fig.1i-red arrows; Fig.S1e-middle panels, red arrows). Knockdown of Cdc42 and MRCKβ in Dbl3-myc expressing ARPE-19 cells confirmed the conserved role of Dbl3 as the apical GEF activating MRCKβ (Fig.1g-i; Fig.S1f). N-WASP plays an important role in actin nucleation and polymerization by activating ARP2/3 in response to binding active Cdc42 (Carvalho et al., 2013; Miki et al., 1998). N-WASP depletion inhibited POS phagocytosis, but not binding, in cells expressing Dbl3-myc (Fig.1g,h). Examination after 30min POS incubation revealed enrichment of both Dbl3-myc and F-actin at the apical cortex in proximity to POS membrane binding sites, which was inhibited by N-WASP siRNA (Fig.1j; Fig.S1g-i). Therefore, RPE phagocytosis requires POS-induced Cdc42 signaling, stimulating two effectors: the myosin-II activator MRCKβ and the regulator of actin polymerization N-WASP.

We next analyzed early morphological changes at POS-membrane contact sites in primary RPE cells. Time course analysis of the POS receptor integrin αvβ5 (Finnemann et al., 1997; Finnemann and Rodriguez-Boulan, 1999; Lin and Clegg, 1998) revealed enriched labelling at POS attachment sites at the onset of POS/RPE contact (10mins) (Fig.S2a). Integrin αvβ5 labelling intensity rapidly increased at attachment sites, peaking at 20min and correlating with an increase in nascent protrusions (Fig.S2a-d). At 30min, there was an increase in protrusions that adopted a cylindrical cup morphology, encircling POS. Therefore, POS contact with the membrane induced αvβ5 integrin clustering at adhesion sites, a common feature of phagocytic receptors (Sobota et al., 2005), and formation of membrane protrusions followed by membrane remodeling into cups around POS before internalization.

Analysis of Cdc42 signaling components at nascent protrusions revealed enrichment in Dbl3, active Cdc42 (Cdc42-GTP), MRCKβ and N-WASP, when compared to non-POS-bound membrane areas (Fig.1k-m; Fig.S3a-f). Cdc42-signaling components continued to localize at mature phagocytic cups (Fig.1k-m). However, at nascent protrusions co-localization with POS was almost complete, whereas at cups co-localization was restricted to the POS-cup interface (Fig.1k,n-q; Fig.S4a-l). Measurement of MLC phosphorylation as an indicator of actomyosin contractility revealed a low activity at nascent protrusions that sharply increased with F-actin enrichment when protrusions transformed to cups (Fig.1r; Fig.S5a-c), indicating a requirement for actomyosin contractility in regulating actin-driven maturation of nascent protrusions to cups to wrap around POS. These results support a role for MRCKβ driven contractility as a regulator of actin dynamics during protrusion maturation into cups.

### MRCKβ/N-WASP control actomyosin dynamics to guide particle wrapping in the RPE

MRCKβ phosphorylates myosin light chain (MLC) and thereby activates myosin-II. Analysis of MLC phosphorylation and F-actin staining revealed maximal phosphorylation after 30 minutes when phagocytic cups had formed and POS were centered within the cups (Fig.2a-c). Both MLC phosphorylation and F-actin enrichment steadily decreased with internalization, which proceeded in two stages: a first one in which POS appeared embedded within the plasma membrane that peaked at 60min and was associated with low but significant MLC phosphorylation, and a second one in which POS was in the cytoplasm and no longer positive for MLC phosphorylation. Inhibition of MRCK activity with BDP5290 abrogated MLC phosphorylation at forming cups and prevented the POS induced increase in total phosphorylated MLC levels (Fig.2d-j; Fig.S6a,b). However, pseudopods still formed but failed to remodel and wrap around POS. No defect in POS binding was observed (Fig.2f; Fig.S6b). Therefore, MRCK activity is not required for pseudopod induction but for remodeling of protrusions to form cups, particle wrapping, and internalization.

**Figure 2.**
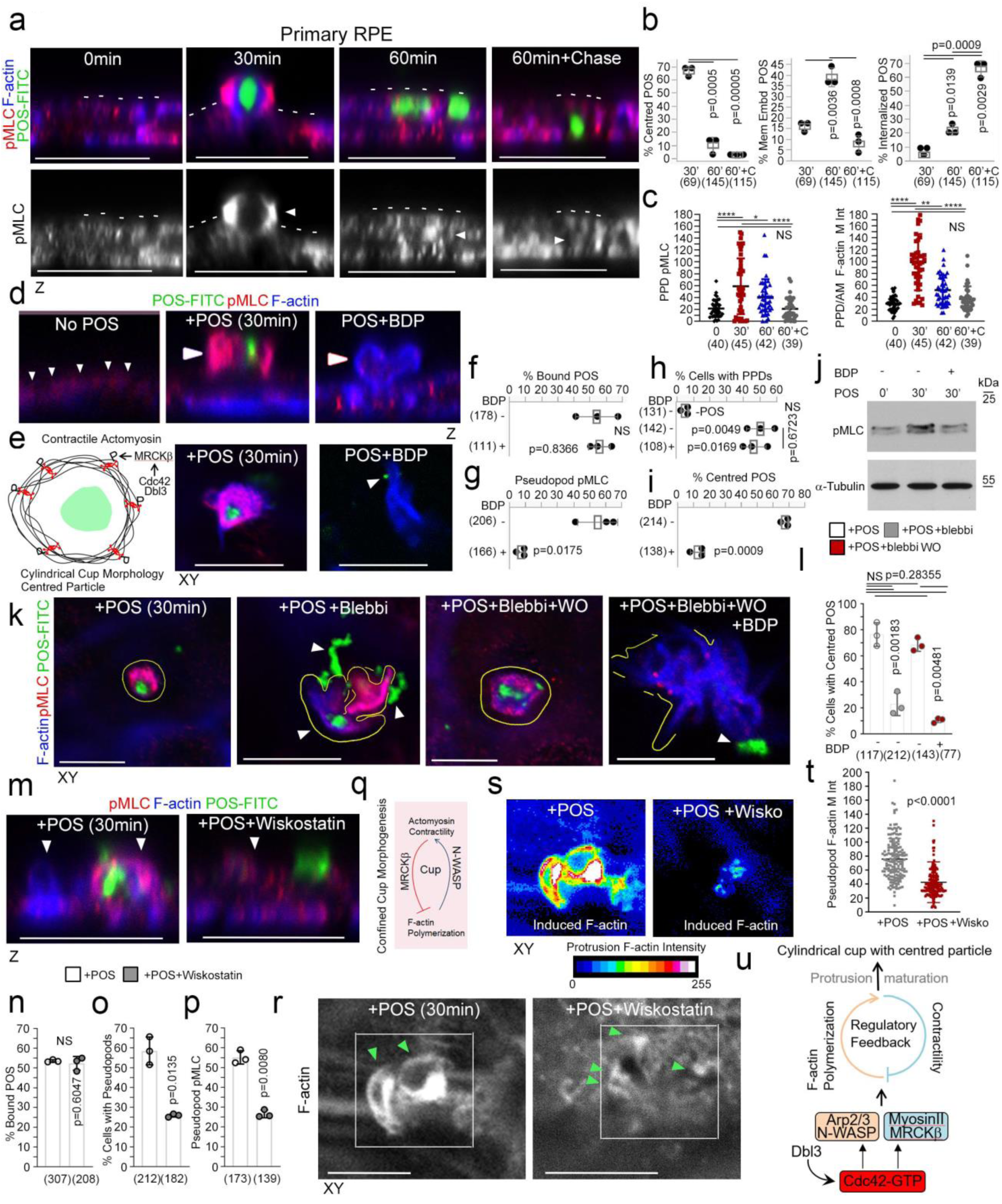
MRCKβ regulates actomyosin dynamics to control POS wrapping in the RPE. **a-c,** Time course of POS-induced cup maturation, closure and POS internalization reveals myosin motor activation and corresponding F-actin enrichment peaks with membrane cups wrapping around POS. **d**-**j**, POS adhesion to porcine primary RPE cultures after 30 minutes incubation results in mature cups wrapping around POS that are enriched with pMLC, whilst MRCKβ inhibition prevented myosin activation and cup remodeling around POS. **k-l**, inhibition of myosin-II motor activity using blebbistatin results in similar membrane remodeling and POS centering defects to MRCKβ inhibition that was reversed within 2h following blebbistatin washout only when MRCKβ was active. **m-p, r-t**, Inhibition of N-WASP activity attenuates protrusion induction and cup maturation with inhibited pMLC. **q,** Schematic figure illustrates a model in which F-actin polymerization mediated at least in part by N-WASP at cups is required for MRCKβ-dependent myosin-II activity, which in turn regulates F-actin polymerization. **u,** Schematic figure illustrates a model in which Dbl3 activates Cdc42 at POS-membrane adhesions to drive a regulatory feedback mechanism between N-WASP and MRCKβ that controls size and shape and wrapping as nascent protrusions mature into cups. Protein staining was quantified as mean intensity. Scale bars represent 10μm, unless highlighted otherwise. All quantifications are based on n=3 independent experiments. Shown are the data points, means ±1SD, the total number of cells analyzed for each type of sample across all experiments, and p-values derived from t-tests.

We next asked whether myosin-II activity is indeed required to control cup morphogenesis. Reversible myosin-II inhibition using blebbistatin (Kovacs et al., 2004) resulted in disorganized pseudopods and loss of efficient POS centering, similar to MRCKβ inhibition or knockdown (Fig.2k-l; Fig.S6c-e). Washout of blebbistatin rapidly restored normal cup organization and POS centering within the forming cup in the absence but not in the presence of the MRCK inhibitor. MRCK activity was also required for MLC phosphorylation at pseudopods. Thus, MRCK regulates actomyosin dynamics during cup morphogenesis and wrapping of POS by regulating myosin-II activity.

In vitro, F-actin polymerization and the formation of actin network architecture defines myosin motor activity (Reymann et al., 2012). Since N-WASP contributes to protrusion induction, we next determined the relationship between Cdc42-driven N-WASP and MRCKβ signaling. Inhibition of N-WASP using Wiskostatin drastically reduced the number of cells forming normal cups (Fig.2m-t), whilst POS binding to the apical membrane of RPE cells was unaffected. Reduced levels of protrusion induction were still observed, possibly reflecting actin polymerization stimulated by the previously described integrin αvβ5-Rac pathway (Mao and Finnemann, 2015) and/or other mutually redundant actin polymerization pathways. These protrusions contained less F-actin (approx. 50%) and less pMLC staining when compared to control cells and did not acquire full cup morphology, indicating that N-WASP is required for full protrusion and cup induction, as well as myosin-II activity (Fig.2p-t). Thus, N-WASP-stimulated F-actin polymerization is required for localized myosin-II activation and phagocytic cup formation (Fig.2q). As myosin-II activity then regulates actin dynamics, protrusion to cup morphogenesis is determined by tightly controlled regulatory feedback between actin polymerization and MRCKβ stimulated myosin-II activation (Fig. 2u).

### MRCKβ drives FAK activity and recruitment of force transducers

We next investigated loss of MRCK activity at an advanced stage of phagocytosis when cup closure and particle internalization are completing. Primary RPE cells were exposed to POS-FITC for 1h followed by a 2h chase in the presence or absence of the MRCK inhibitor BDP5290; alternatively, MRCKβ was knocked down by RNA interference (Unbekandt et al., 2014; Zihni et al., 2017b). Analysis of confocal z-sections revealed that both methods of MRCKβ inactivation inhibited POS internalization (Fig.3a,b,e; Fig.S7b,g). Electron microscopy of RPE cells fed with GOLD-labelled instead of fluorescent POS confirmed inhibition of phagocytosis (Fig.3b; Fig.S7j). As in shorter time courses, MRCKβ was not required for POS attachment to the membrane, indicating integrin αvβ5-POS binding was not affected (Fig.3d; Fig.S7a,f,i).

**Figure 3.**
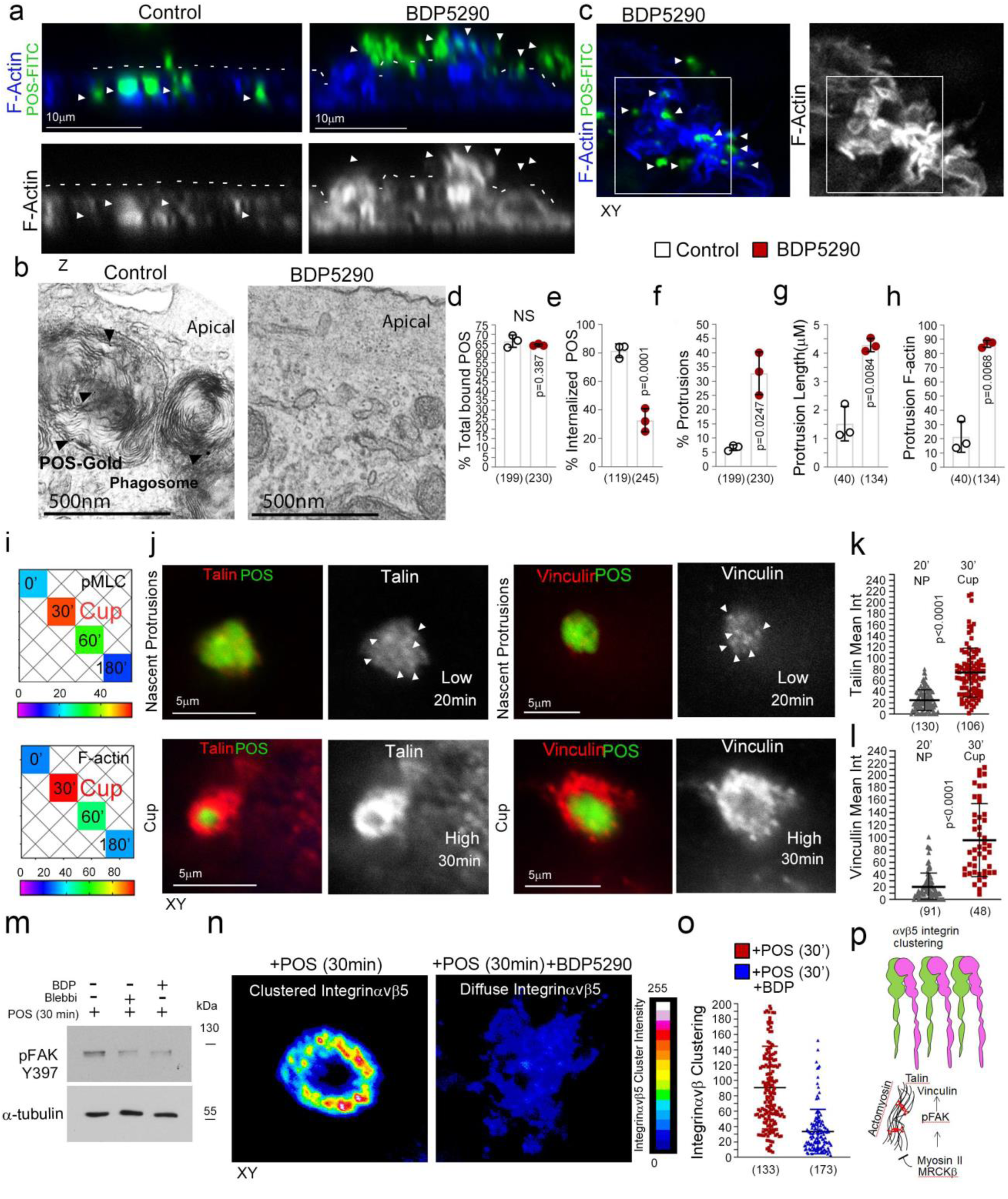
MRCKβ controls mechanotransduction and integrin adhesion. **a-h**, Inhibition of MRCKβ activity in primary porcine RPE cultures results in defective POS-induced apical membrane remodeling with stalled F-actin-based protrusions and inhibited phagocytosis as determined by confocal z-sections of porcine primary RPE cells stained with Atto647-Phalloidin; POS were labelled with FITC. White dashes highlight apical membrane and arrows, internalized POS on the left panels and extracellular POS on the right panels. Transmission electron microscopy of primary porcine RPE cells using GOLD-labelled POS reveals MRCK kinase inhibition using BDP5290 results in an inhibition of phagocytosis. The quantifications in d-h are based on n=3 independent experiments and show data points, means ±1SD (top of bar), the total number of cells. **i,** Heat map of POS-RPE incubation time course. Note, both pMLC and F-actin labelling peak at cup formation. **j-l,** Confocal microscopy analysis of primary RPE treated with POS-FITC revealed recruitment of mechanosensors talin and vinculin at maturing cups, concomitant with robust actin polymerization and increased actomyosin contractility. Values were calculated by subtracting mean staining intensity of apical membrane that is not attached to POS, in the same population **m**, Inhibition of actomyosin contractility, during cup maturation, by inhibiting myosin-II motor activity using blebbistatin or MRCKβ kinase activity using BDP inhibits FAK phosphorylation. **n**,**o,** Inhibition of MRCK-driven actomyosin contractility inhibits clustering of αvβ5 integrin at cups. Note, αvβ5 integrin appears diffuse at cups along with loss of cup morphology and POS centering. Protein staining was quantified as mean intensity. **p**, Sschematic diagram illustrating the dual function of MRCKβ driven contractility that limits actin polymerization to control maturation of protrusions into cups, whist activating FAK to drive molecular clutch assembly. Scale bars represent 10μm, unless highlighted otherwise. The quantifications are based on n=3 independent experiments and show data points, means ±1SD (top of bar), the total number of cells analyzed for each type of sample across all experiments and p-values derived from t-tests.

MRCKβ inactivation resulted in an irregular apical membrane at POS-bound sites with elongated F-actin-positive protrusions (Fig.3a,c,f-h; Fig.S7a-e,i). Since MRCKβ activates myosin-II, the defect in pseudopod remodeling during normal cup closure is in agreement with a function of myosin-II-dependent contractility in limiting actin polymerization by promoting depolymerization, as demonstrated with *in vitro* models (Reymann et al., 2012). The prolonged pseudopods failed to encircle POS, appearing irregular with misaligned particles in agreement with the requirement for MRCKβ-activated contractility for POS engulfment to form mature cups (Fig.3a,c; Fig.S7a,b,i). Thus, phagocytosis requires a regulatory feedback mechanism between actin polymerization and MRCKβ-activated myosin-II motors to control transformation of POS-adhesion-induced protrusions into phagocytic cups.

Protrusions transform into mature cups with increased F-actin accumulation and myosin-II activity (Fig.3i), indicating increased mechanical force may act on the phagocytic cup to drive cup closure and internalization. Strikingly, there was a progressive increase in the recruitment of talin and vinculin as protrusions matured into cups, correlating with F-actin and pMLC accumulation (Fig.3j-l). Vinculin and talin are recruited to integrin complexes in response to cytoskeletal tension and form a molecular clutch that transduces mechanical force to integrins, a function regulated by FAK at basal focal adhesions of spread cells (Chen, 2008; Oria et al., 2017) Inhibition of MRCKβ or myosin-II inhibited FAK-Y397 phosphorylation in the presence of POS, which occurs after 30 minutes in control cells and correlates with cup formation (Fig. 3m). Thus, MRCKβ-stimulated myosin-II activation stimulates FAK activation and recruitment of mechanical force transducers at POS adhesion sites.

Cytoskeletal tension acting on receptors induces clustering to strengthen particle adhesion (Oria et al., 2017; Sobota et al., 2005). Inhibition of MRCKβ activity did not affect initial accumulation of αvβ5 integrins at nascent protrusions in agreement with the low level of actomyosin contractility observed at this early stage (Fig.S7k,l). However, integrin clustering at mature cups was inhibited by MRCK inhibition (Fig.3n,o; Fig.S7m). Therefore, increasing MRCKβ-stimulated actomyosin activity drives phagocytic cup maturation and leads to the activation of mechanosensitive signaling and integrin clustering prior to internalization (Fig.3p).

### MRCKβ is required for RPE function *in vivo*

We next asked whether MRCKβ is required for retinal function in vivo. To test this, we deleted MRCKβ in the RPE of adult mice, a non-proliferative polarized epithelium, by subretinal injection of lentiviral CRISPR-Cas9 vectors, a method that enables specific transduction of the RPE (Fig.4a-b; Fig.S8a,b) (Georgiadis et al., 2010). Analysis of retinal tissue sections by confocal microscopy revealed that mice maintained for 21days post-injection displayed a striking thinning of the outer nuclear layer (ONL), a hallmark of advanced retinal degeneration (Fig.S8c) (Edwards and Szamier, 1977). At 7 days post-injection, the ONL thickness and outer limiting membrane F-actin intensity were unchanged (Fig.4c,d), indicating that the general architecture and integrity of the retina were maintained. However, MRCKβ knockout RPE cells displayed a reduced apical-basal F-actin intensity ratio (Fig.4c,d), indicating that the apical membrane-associated cytoskeleton was already affected. A second lentiviral CRISPR-Cas9 vector that targeted a different region of the MRCKβ gene, generated a similar effect on apical-basal F-actin distribution (Fig.S8e-g). The apical RPE membrane in vivo is covered by extended microvilli that function in phagocytosis (Bonilha et al., 2006); hence, we investigated the ultrastructure of the RPE using transmission electron microscopy (TEM). In control RPE cells, apical microvilli were ordered and packed into linear arrays that extended to and surrounded POS as previously reported (Fig.4,h,i,k) (Bonilha et al., 2006). RPE cells contained internalized phagosomes indicating functional RPE (Fig.4h,i). In contrast, microvilli of MRCKβ-knockout RPE in contact with POS appeared disordered and more spread out (Fig.4j,l). Measurement of microvilli thickness and packing into sheets revealed a decrease in microvilli packing order (Fig.4m,n; Fig.S4d). Strikingly, MRCKβ-targeted RPE cells contained >7 times less cytoplasmic phagosomes, which was paralleled by an accumulation of extracellular shed POS (Fig.4e-g). Thus, MRCKβ is required for RPE phagocytosis and retinal integrity *in vivo*.

**Figure 4.**
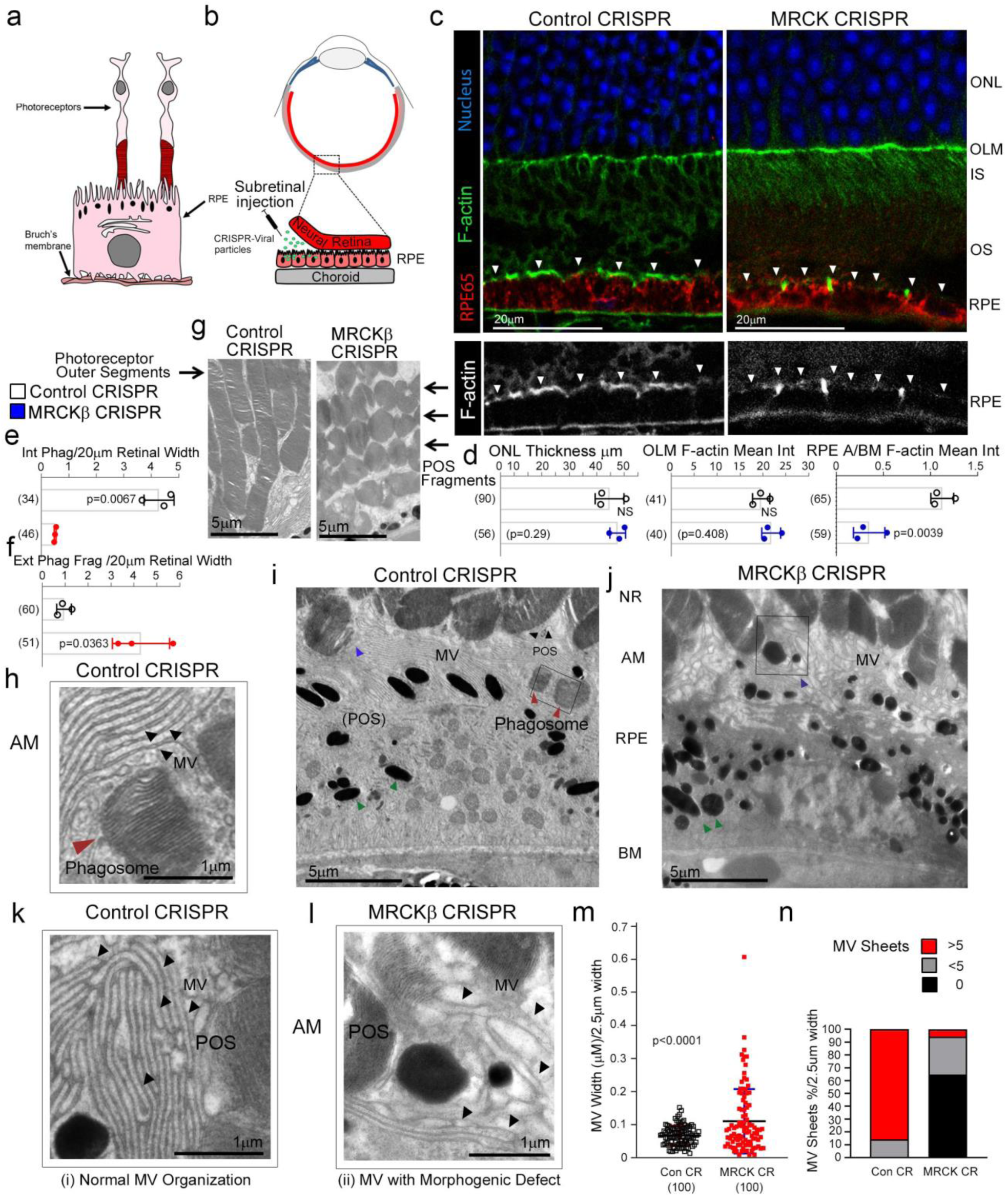
MRCKβ is required for RPE function and retinal integrity in vivo. **a**, Schematic illustration of RPE and photoreceptor architecture from the retina. Note, photoreceptor outer segments (POS) are in contact with microvilli at the apical membrane of RPE. As POS is shed at the tips the resultant fragments expose membrane components that initiate the phagocytic cycle. **b**, Schematic illustration of subretinal delivery of lentiviral vectors. Note, injection of viral particles into the subretinal space results in specific transduction of the RPE layer. **c**, **d**, Confocal immunofluorescence analysis of retinal sections from mice injected with control or MRCKβ knockout vectors reveal no significant difference in the overall retinal structure after 7 days but reduced apical F-actin staining, with reference to basal F-actin, in the MRCKβ−deficient RPE. White arrows highlight apical F-actin cortex. **e**-**j**, Transmission electron microscopic analysis showing that RPE from control CRISPR-vector-injected mice internalized POS fragments whereas internalization in MRCKβ CRISPR-vector injected mice was inhibited, resulting in accumulation in the extracellular space. Red and blue arrows highlight internalized and internalizing POS fragments, respectively, in control RPE. Green arrows highlight melanosomes in both control and MRCKβ knock-out samples. **k-n**, Transmission electron microscopic analysis of RPE apical microvilli architecture in control and MRCKβ CRISPR-vector injected mice. Note, the quantifications show that microvilli from controls display ordered sheet packing and individual morphology whereas from MRCKβ knockout animals a disordered appearance of microvilli with a loss of the sheet arrangement and irregular individual thickness. Black arrows highlight individual mv strands. Quantifications show means ± 1SD, n=3, and p-values derived from t-tests.

### MRCKβ-signaling regulates phagocytosis in Macrophages

We next tested the involvement of MRCKβ in FcR-mediated macrophage phagocytosis, which is Cdc42-dependent (Caron and Hall, 1998; Massol et al., 1998). In macrophages differentiated from THP-1 cells, MRCKβ associated with the cell cortex (Fig.5a). Fluorescent, IgG-opsonized Zymosan-yeast particles were efficiently internalized by control siRNA-treated cells, as determined by confocal z-section analysis, but not cells treated with siRNAs targeting MRCKβ. Knockdown led to increased accumulation of opsonized particles embedded in the plasma membrane cortex, indicating stalled cup closure and attenuated full internalization (Fig.5b-f). Myosin-II, the substrate of MRCKβ (Unbekandt et al., 2014; Zhao and Manser, 2015; Zihni et al., 2017b), is required for completion of cup closure during FcR-mediated phagocytosis (Barger et al., 2020), and myosin-II inactivation leads to a similar phenotype as displayed by MRCKβ-depleted cells (Barger et al., 2020; Clarke et al., 2010). Hence, our results indicate that MRCKβ-stimulated actomyosin contractility regulates completion of phagocytic cup closure and internalization in RPE and macrophages. The distinct employment of Cdc42/MRCKβ signaling of different phagocytic machineries in the RPE and macrophages indicates plasticity of the signaling module, enabling the Cdc42/MRCKβ mechanism to be adapted to the requirements of different phagocytic receptors (Fig.5g).

**Figure 5.**
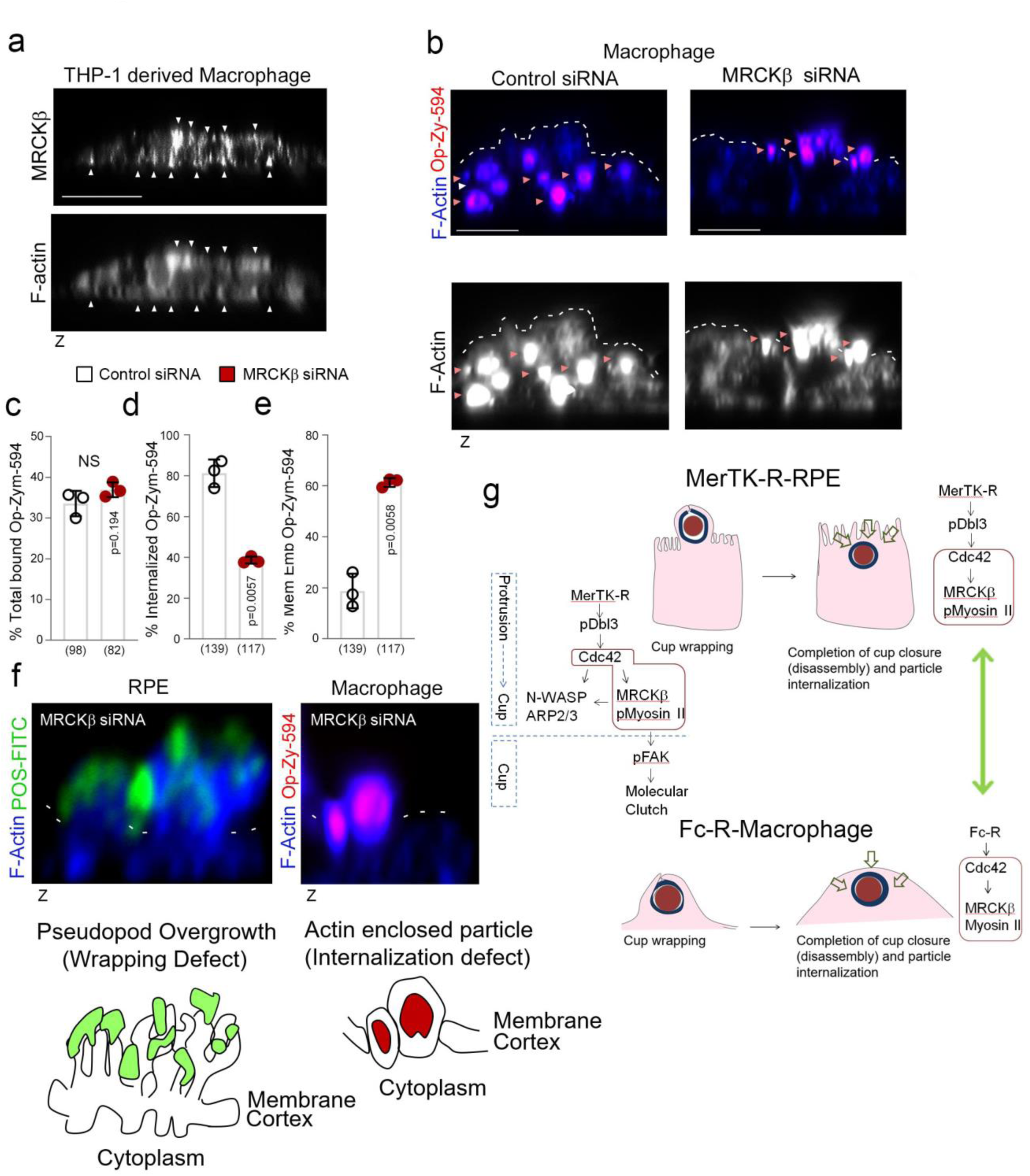
MRCKβ drives engulfment in Cdc42-mediated FcR-macrophages. **a**, Confocal z-sectional analysis of THP-1 derived macrophages reveal MRCKβ localizes along the actin cortex. **b-e,** siRNA-mediated knockdown of MRCKβ in THP-1-derived macrophages demonstrates its requirement for Fc-receptor-mediated IgG-opsonized particle internalization. Note that forming phagosomes are embedded in the cortex. **f,** Comparison of membrane defects and inhibition of particle internalization in MerTK-mediated RPE and Fc-Receptor-mediated macrophage phagocytosis following MRCKβ knockdown. Note, loss of MRCKβ signaling in the RPE results in F-actin overgrowth and disordered pseudopods that do not wrap around particles whereas in macrophages the particle is partially embedded in the cortex and enclosed by F-actin. In both cell types, particles were added for long time points allowing particle internalization in control cells. **g,** Schematic model of proposed role of Cdc42/MRCKβ-signaling in MerTK-receptor-mediated RPE and Fc-receptor-mediated macrophage phagocytosis. In RPE cells, Cdc42/MRCKβ-signaling cooperates with N-WASP to regulate actin dynamics from the onset of protrusion induction to guide cup maturation. In cups, Cdc42/MRCKβ-signaling activates FAK to drive molecular clutch assembly and receptor clustering to control wrapping of POS. In both RPE and macrophages, MRCKβ is required for actomyosin contractility-driven particle internalization. Scale bars represent 10μm. The quantifications are based on n=3 independent experiments and show data points, means ±1SD, the total number of cells analyzed for each type of sample across all experiments and p-values derived from t-tests.

### MerTK activates Dbl3 to drive phagocytosis in the RPE

We next asked whether the upstream GEF of MRCKβ, Dbl3, is activated by MerTK to stimulate phagocytosis. As αvβ5 integrin and the Dbl3-activated Cdc42 signaling machinery, MerTK was enriched at nascent protrusions and almost completely co-localized with POS (Fig.6a-d; Fig.S9a-c). In mature cups, integrin αvβ5 only significantly co-localized with POS at POS-membrane contacts, similar to the Cdc42 signaling components, whereas MerTK still displayed significant co-localization (Fig.6c,d; Fig.S9b-c). Higher co-localization of MerTK with POS was expected as POS binding promotes cleavage of the extracellular N-terminal domain that then quenches still available MerTK binding sites on POS (Law et al., 2015). Activated tyrosine kinase pSrc Y416, known to be activated by MerTK (Shelby et al., 2015), also localized to POS-induced protrusions (Fig.6b; Fig.S9d).

To test whether Dbl3-activated actomyosin dynamics in response to POS requires MerTK activation, primary porcine RPE cells were cultured in serum-free medium to eliminate effects of serum MerTK ligands. The cells were then stimulated with POS-FITC in the absence or presence of Gas-6, the ligand that links MerTK to POS (Etienne-Manneville and Hall, 2002; Hall et al., 2002; Kevany and Palczewski, 2010), without or with N-WASP or MRCK inhibitors. POS binding to the apical membrane was observed without addition of Gas-6, reflecting binding to the integrin, but induction of protrusions was inhibited (Fig.6e-left panel, f; Fig.S9e-left panel, g). Addition of Gas-6 together with POS induced an increase in protrusions positive for pMLC (Fig. 6e-right panel, f; Fig.S9e-top middle panel). Inhibition of N-WASP prevented protrusion formation, and inhibition of MRCKβ prevented pMLC induction and cup-POS centering (Fig.6f,g,h; Fig.S9e-bottom panels). Knockdown of Dbl3 attenuated pMLC-positive cups and POS wrapping, supporting a role for the GEF downstream of MerTK and upstream of MRCKβ (Fig.6i,j; Fig.S9e-top right panel, f).

**Figure 6.**
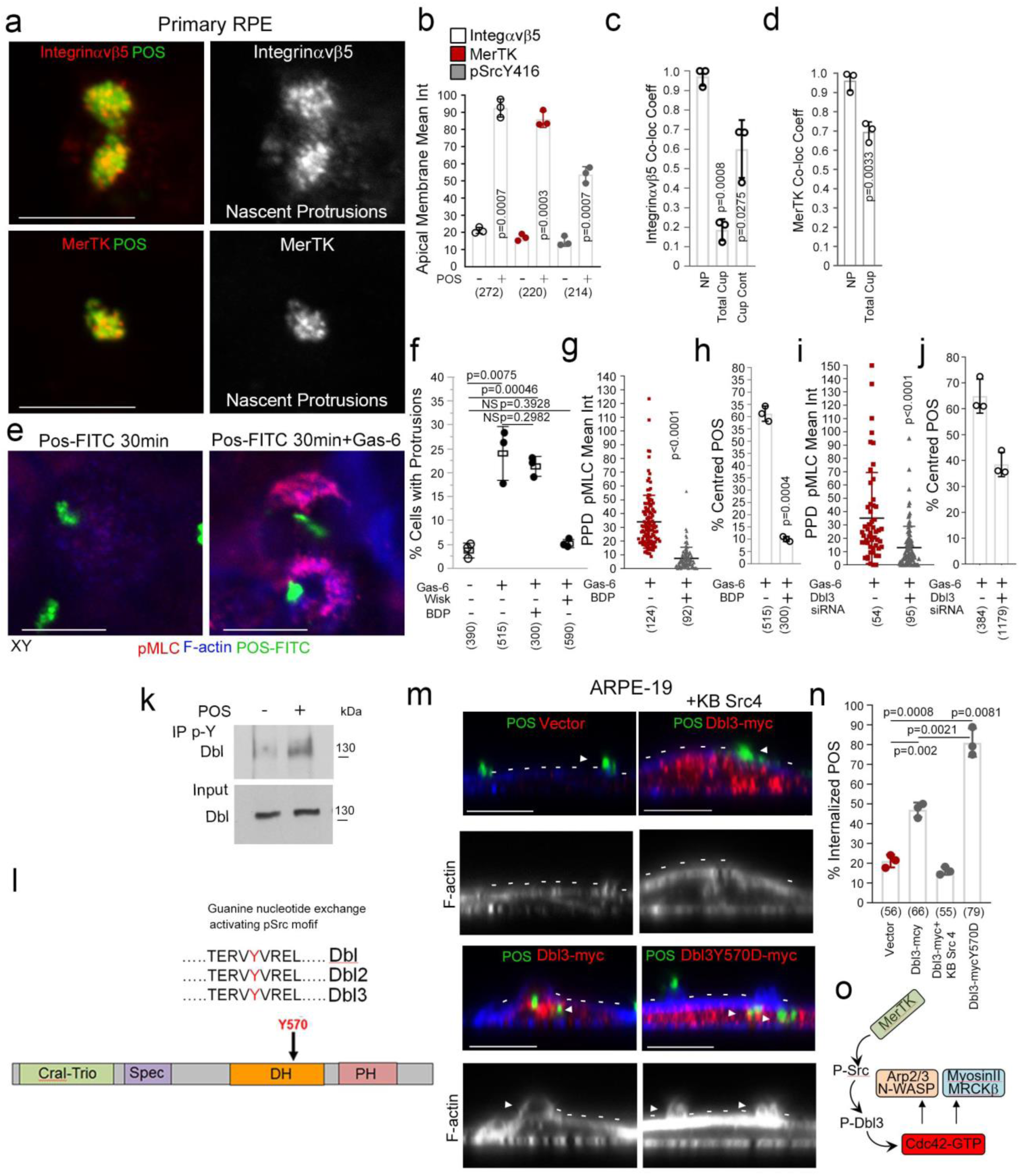
MerTK stimulates Dbl3-activated actomyosin dynamics. **a**-**d**, Integrin αvβ5, MerTK tyrosine kinase receptor and its effector Src are enriched at and co-localize with POS at nascent protrusions in primary porcine RPE cells. **e,h,** stimulating primary porcine RPE cells with POS in serum-deprived conditions does not induce protrusions unless ligand for MerTK, Gas-6, is added. Gas-6 promotes MRCK-dependent pMLC activity at cups and POS centering. **i,j,** Dbl3 signaling is required for POS-Gas-6 stimulated pMLC activity at cups and POS centering. **k**, Immuno-blot showing immunoprecipitation of Dbl by anti-phosphotyrosine antibody following POS stimulation of primary porcine RPE cells. **l,** Schematic illustration of a predicted Src/tyrosine phosphorylation motif in the Dbl3 DH domain. **m,n,** Exogenous expression of Dbl3-myc wild type in the presence of a c-Src inhibitor blocks protrusion induction and POS engulfment in ARPE-19 cells, whereas Dbl3Y570D-myc stimulates increased POS internalization. Arrows in F-actin images highlight protrusion and cup induction by Dbl3-myc. **o,** Schematic illustration of proposed model in which phosphorylation of Dbl3 following MerTK stimulation of Src activates Cdc42 signaling. Scale bars represent 10μm. All quantifications are based on n=3 independent experiments and show the data points, means ±1SD, the total number of cells analyzed for each type of sample across all experiments, and p-values derived from t-tests.

Localization of the MerTK effector c-Src at nascent protrusions along with MerTK and Dbl3 suggests that Dbl3 activation my involve tyrosine phosphorylation. Immunoprecipitation of phospho-tyrosine containing proteins in primary porcine RPE cells stimulated with POS for 20 minutes revealed tyrosine phosphorylation of Dbl3 (Fig.6k). Bioinformatics analysis using NetPhos3.1 suggested a conserved tyrosine phosphorylation motif TERVYVREL in the Dbl-homology (DH) domain that carries GEF activity (Fig.6l; Fig.S9h). This motif had been shown to be phosphorylated in Dbl1, a functionally distinct splice variant of Dbl3 that lacks apical targeting information but contains a homologous DH domain (Gupta et al., 2014; Zihni et al., 2014). The TERVYVREL motif of Dbl1 is a Src family kinase target, and phosphorylation activates its GEF activity *in vitro* (Gupta et al., 2014). Src tyrosine kinase inhibition blocked Dbl3-myc driven phagocytosis in ARPE19 cells (Fig.6m,n). Conversely, expression of a phosphomimetic Dbl3Y570D mutant increased internalization. Thus, Dbl3-activation of MRCKβ and subsequent control of phagocytosis requires Src activity (Fig.6o).

### Dbl3Y570D recuses phagocytosis in MerTK-deficient RPE

Dbl3 activates MRCKβ and its co-effector N-WASP to drive phagocytosis following POS-MerTK ligation; hence, enhancing Dbl3-signaling may be sufficient to stimulate POS internalization in phagocytosis-deficient RPE. To test this, we employed RPE cells generated from induced pluripotent stem cells (iPSC) derived from a retinitis pigmentosa individual carrying biallelic nonsense mutations in the MerTK gene, leading to loss of MerTK protein expression, phagocytosis deficiency and blindness (Fig.7a-d; Fig.S10a,b) (Ramsden et al., 2017). MerTK-deficient iPSC-derived RPE cells differentiate and polarize normally with functional integrin αvβ5 receptors, except for the absence of MerTK protein (Ramsden et al., 2017). As previously reported, these cells bound POS efficiently but displayed poor phagocytosis (∼11%) (Fig.7e-g). Endogenous Dbl3 co-localized with attached POS characteristic of early adhesion sites with no significant protrusion or cup formation detected, indicating lack of Dbl3-induced signaling in agreement with a model in which MerTK signaling activates Dbl3 (Fig.7h,i; Fig.S10c,f,g). Expression of Dbl3-myc in MerTK-deficient RPE only weakly increased phagocytosis (∼22%) (Fig.7e,g, k-middle panel; Fig.S10d,h,i). In contrast, expression of the phosphomimetic Dbl3Y570D mutant increased phagocytosis to 87.0% and promoted efficient maturation of POS adhesion sites to phagocytic cups with characteristic co-localization of Dbl3 with POS at their inner surface (Fig;7e,g,h,j-k-right panel; Fig.S10e,j,k,l,). Thus, Dbl3Y570D expression functionally mimics MerTK signaling and rescues phagocytosis in MerTK-deficient RPE cells (Fig.8). Stimulation of the MRCKβ pathway by Dbl3 is therefore sufficient to rescue phagocytosis in MerTK-deficient cells, indicating that it represents the main phagocytic effector pathway of MerTK.

**Figure 7.**
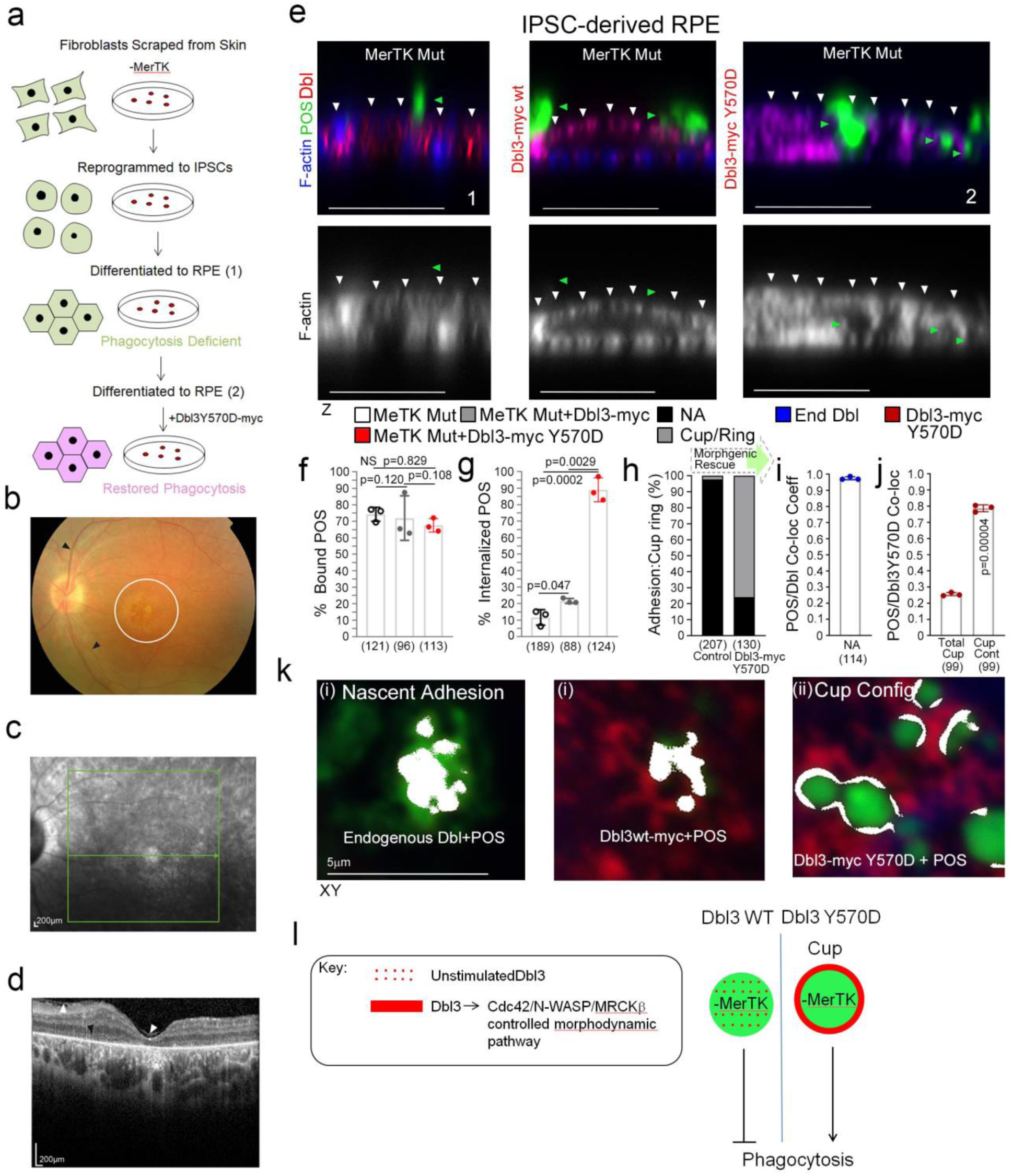
Dbl3Y570D rescues phagocytosis in mutant MerTK iPSC-derived RPE cells. **a**, Schematic outline of RPE cell generation from the MerTK mutant fibroblasts. Note, fibroblasts were derived from a blind individual suffering from retinitis pigmentosa (RP). **b,** Fundus image taken from a MerTK-deficient RP individual’s right eye showing the retinal vessels (black arrows) and macula (white circle). **c**, Shows a red free image of the macula with the green arrow representing the position of the line scan in **b**. **d**, is a spectral domain optical coherence tomography image of the left fundus, through the fovea, showing a thinning of the retina at the fovea (normally over 200µm) and a loss of retinal lamination, especially centrally (total loss of ellipsoid zone, black arrowhead), the white arrowhead shows an epiretinal membrane. **e-g**, Confocal immunofluorescence z-section analysis of MerTK mutant RPE transfected with wild type Dbl3 or Dbl3Y570D and exposed to POS-FITC. Note, RPE cells expressing Dbl3Y570D undergo efficient recovery of phagocytosis. **h-k,** Analysis of POS adhesion sites to cup population ratios in mutant cells and cells expressing Dbl3Y570D. Note, co-localization analysis by Pearson’s coefficient calculation reveals that MerTK-deficient RPE cells contain almost exclusively Dbl rich adhesion sites that fail to transform into protrusions depicted in white, whereas mutant cells expressing Dbl3Y570D undergo cup maturation with characteristic Dbl-POS colocalization at contact sites depicted as white rings and point contacts. Scale bars represent 10μm, unless highlighted otherwise. All quantifications are based on n=3 independent experiments and show the data points, means ±1SD;, the total number of cells analyzed for each type of sample across all experiments, and p-values derived from t-tests are indicated. **l,** Schematic illustrating efficient repair of nascent adhesion to cup maturation by Dbl3Y570D signaling in MerTK-deficient iPSC-derived RPE.

**Figure 8.**
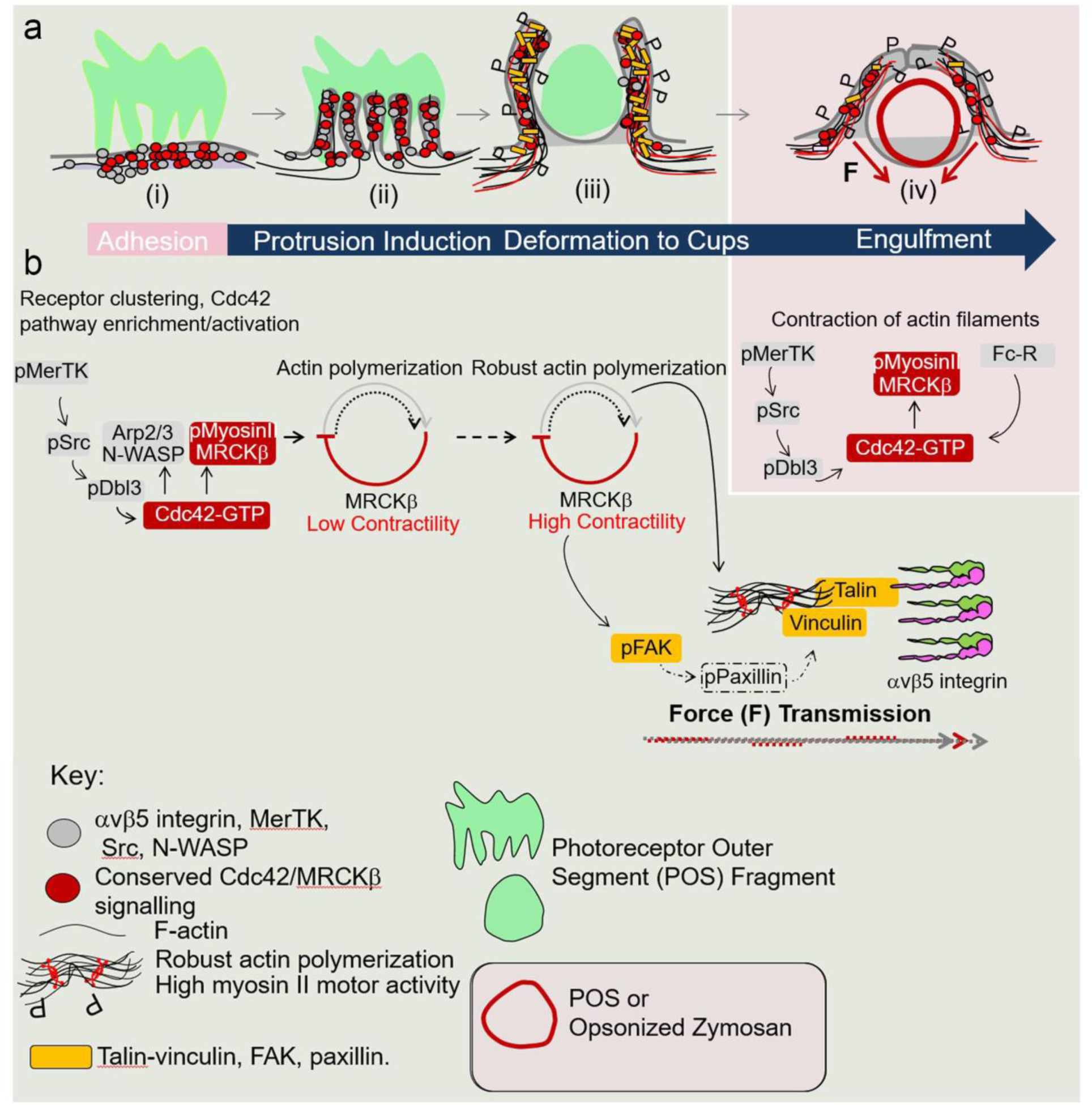
Proposed model of MRCKβ signaling in distinct phagocytes. Schematic illustration of proposed model of MRCKβ-controlled morphodynamic signaling in MerTK receptor mediated retinal pigment epithelial cells (green background and pink background) and Fc-Receptor mediated macrophage signaling (pink background). **a**,**b** In the RPE, MerTK-Cdc42-driven MRCKβ signaling progresses through distinct phases with a lower activity during initial pseudopod induction and high activity, concomitant with increased F-actin, during deformation of protrusions to mediate particle wrapping. Continued MRCKβ-driven myosin II activity stimulates FAK activation of paxillin (dashed outline represents a putative link based on the established mechanosensory bridge in migratory cells), recruitment of mechanotransducers vinculin and talin, and receptor clustering at cups to coordinate particle wrapping. In both RPE and macrophages, MRCKβ signaling is required for the completion of cup closure and internalization.

## DISCUSSION

This study identifies an MRCKβ-driven signaling mechanism that spatiotemporally regulates actomyosin contractility to guide distinct morphological membrane transformations required for RPE and macrophage phagocytosis. In the RPE, this mechanism functions downstream of MerTK, a common receptor generally found on phagocytes that internalize dead cells and cell remnants such as POS in the retina. Expression of a mutant form of the Cdc42 exchange factor Dbl3 that mimics MerTK-activation is sufficient to rescue phagocytosis in MerTK-deficient RPE cells, indicating that it represents the principal phagocytic effector pathway activated by MerTK.

In the RPE, MRCKβ is activated by MerTK-stimulated phosphorylation of the apical Cdc42 GEF Dbl3 to drive myosin-II-mediated actomyosin contractility to guide maturation of particle-induced protrusions into cups in cooperation with N-WASP. POS-membrane adhesion results in rapid enrichment of the receptors αvβ5 integrin and MerTK at adhesion sites along with the Cdc42 signaling machinery, resulting in localized MerTK-activated Cdc42 signaling. This stimulates actin polymerization, at least in part due to N-WASP, which forms the basis for generating membrane protrusions. MRCKβ-stimulated contractility then controls maturation of protrusions into cups and particle wrapping by regulating actomyosin dynamics. Inactivation of MRCKβ results in extended growth of pseudopods, indicating that MRCKβ-regulated contractility is required to limit actin polymerization. These findings are in agreement with a model in which the interdependence of actin polymerization and contractility creates the conditions for the spatial and temporal control of actin cytoskeletal dynamics required to drive complex morphodynamic processes (Reymann et al., 2012).

MRCKβ mediated myosin-II activity proceeds to guide particle engulfment and contractile force-dependent pulling into the cell by coupling to additional cytoskeletal mechanisms. During the transition from particle wrapping to engulfment, MRCKβ-activated contractility promotes formation of a force-transducing bridge between integrins and accumulating actin fibers by activation of FAK, stimulating recruitment of vinculin and talin, as well as integrin clustering.

αvβ5 integrin has been shown to activate FAK at an earlier step, on initial binding to POS, which has been proposed to contribute to the rhythmic burst activity of phagocytosis by RPE in vivo by enhancing MerTK activity (Finnemann, 2003; Finnemann et al., 1997). Rac1 is activated by αvβ5 integrin, although its precise role in phagocytosis is unclear since its activity is independent of FAK and MerTK (Mao and Finnemann, 2012). Rac1 may contribute to the remaining protrusion formation observed in N-WASP depleted cells, indicating cooperative mechanisms driving actin polymerization. Cdc42 and Rac 1 have been shown to display spatiotemporally coordinated localization and morphogenic transformation profiles in Fc-R mediated macrophage engulfment (Hoppe and Swanson, 2004); thus, future work to determine the mechanistic relationship between the two GTPases is likely to provide important information about coordination of their functions in phagocytosis. Since FAK activation by αvβ5 integrin first enhances MerTK activity and FAK activity stimulated by MRCKβ then functions later most likely to promote recruitment of mechanotransducers and integrin clustering, our results suggest a biphasic activation of FAK. FAK activation by MRCKβ is reminiscent of mesenchymal migration, where myosin-II-dependent FAK-mediated phosphorylation of paxillin promotes vinculin binding to talin and actin to reinforce the force transducing link between the cytoskeleton and the ECM (Chen, 2008; del Rio et al., 2009; Humphries et al., 2007; Pasapera et al., 2010; Thievessen et al., 2013). Since spreading of migratory cells on a matrix is topologically similar to phagocytosis of particles (Grinnell, 1984), our results support a model that postulates the existence of signaling mechanisms that control the actomyosin cytoskeleton during phagocytosis in a similar manner as during cell spreading. Complement receptor-mediated macrophage phagocytosis, which is myosin-II-independent, also stimulates recruitment of such ‘molecular clutch’ proteins, suggesting that force-generation during cup morphogenesis in macrophages can be based on distinct mechanisms (Case and Waterman, 2015) Jaumouille et al., 2019)

In FcR-mediated macrophage phagocytosis, particle binding induces a distinct step-specific pattern of Cdc42 activation, which is required for myosin-II-dependent completion of cup closure and internalization (Hoppe and Swanson, 2004). Our data now indicate that Cdc42/MRCKβ signaling is responsible for myosin-II activation during completion of cup closure and internalization. Thus, while Cdc42/MRCKβ morphogenetic signaling is conserved downstream of different phagocytic receptors, it is already required at earlier steps during MerTK-regulated phagocytosis in the RPE, indicating that plasticity of the signaling machinery enables adaptation to the specific requirements of different phagocytic receptors.

The essential function of MRCKβ-controlled morphogenetic signaling in distinct receptor-mediated phagocytes and its importance for retinal integrity indicate that its associated mechanism may have a use as a therapeutic target in disease. As Dbl3 is activated by POS binding to MerTK, receptor-independent activation of Dbl3 signaling should stimulate phagocytosis. Indeed, expression of phosphomimetic Dbl3 in retinitis pigmentosa iPSC-derived MerTK-deficient RPE cells rescued phagocytosis, indicating that it represents the main phagocytic effector pathway downstream of MerTK. The ability to rescue phagocytosis despite loss of phagocytic receptor function may lead to broader therapeutic applications in the RPE beyond MerTK dysfunction, including age-related decline in phagocytosis, which may contribute to age-related macular degeneration, a leading cause of blindness (Inana et al., 2018; Sparrow et al., 2010). MerTK-regulated phagocytosis is important for the function of diverse phagocytes, and its deregulation is thought to contribute to difficult to treat diseases such as neurodegeneration (Brosius Lutz et al., 2017) (Myers et al., 2019) (Haukedal and Freude, 2019). Similarly, macrophages and other phagocytic cells of the immune system play important roles in infection, chronic inflammatory disease and cancer, and can promote tissue repair as well as damage. Therefore, understanding the molecular mechanisms that control phagocytosis and identifying new approaches to modify such mechanisms will contribute to designing innovative new therapies for a wide range of diseases (Ardura et al., 2019).

## Materials and METHODS

### Cell culture and generation of mammalian expression and viral vectors

Human ARPE-19 cells were obtained from ATCC. Primary porcine RPE cells were isolated from pig eyes as described previously(Tsapara et al., 2010). ARPE-19 and porcine RPE cells were grown in DMEM supplemented with 10% fetal bovine serum. Once isolated porcine RPE were confluent they were cultured in DMEM medium containing 1% FBS and passaged no more than 2 times. For phagocytosis assays, the cells were transferred to medium containing 2.5% fetal bovine serum. Fresh batches of ARPE-19 cells from a contamination-free stock that had been tested for mycoplasma (MycoAlert; Promega Inc.) were used every 6 to 8 weeks. Cells were then weekly stained with Hoechst dye to reveal nuclei and DNA of contaminants such as mycoplasma. iPSC cells carrying inactivating mutations in MerTK were characterized previously and were differentiated into RPE cells as described (Ramsden et al., 2017). MRCK inhibitor BDP5290 was generously provided by Michael Olson (Cancer Research UK Beatson Institute, Glasgow, UK) and was synthesized by the Cancer Research UK Beatson Institute Drug Discovery Group. It was used at a concentration of 10mM. Blebbistatin (used at 10µM) and Wiskostatin (used at 5µM) were purchased from Tocris Bioscience. The expression plasmid for Dbl3-myc was as described (Zihni et al., 2014). Dbl3Y570D substitution was introduced into Dbl3-myc by PCR using the CloneAmp HiFi PCR premix and In-Fusion cloning (Clontech Inc.). Constructs were verified by sequencing. CRISPR Cas9 lentiviral vectors were obtained from Sigma-Aldrich and were constructed in pLV-U6g-EPCG. The sequences targeted were for MRCKβ 5’-CCTGGACGGGCCGTGGCGCAAC-3’ and 5’-ACACCGAGTGCAGCCAC TCGG-3’, and for Dbl3 5’-TGGAGATCGAAGACTGGATACATGG-3’. Plasmids were transfected using TransIT-X2 (Cambridge Bioscience)

### Transfection

Cells were cultured and transfected with Lipofectamine RNAiMAX (Thermo Fischer Scientific) or Vitromer Blue (Lipocalyx) using siRNAs targeting the following sequences: human N-WASP 5’-CAGAUACGACAGGGUAUCCAA-3’ and 5’-UAGAGAGGGUGCUCAGCUAAA-3; porcine Cdc42, GAUGACCCCUCUACUAUUG; porcine Dbl3 5’-AAGACAUCGCCUUCCUGUC-3’ and 5’-AUACCUGGUCUUCUCUCAA-3’; and porcine MRCKβ 5-CGAGAAGACUUCGAAAUAA-3’ and 5’-AGAGAAGACUUUGAAAUAU-3’.

The non-targeting control siRNAs and those for human Cdc42, MPXKβ, and Dbl3 were as described previously (Zihni et al., 2014; Zihni et al., 2017b).

### Maintenance of mice

Wild-type mice (C57BL/6J) were purchased from Harlan Laboratories (Blackthorn, UK). All mice were maintained under cyclic light (12 h light–dark) conditions; cage illumination was 7 foot-candles during the light cycle. All experiments were approved by the local Institutional Animal Care and Use Committees (UCL, London, UK) and conformed to the guidelines on the care and use of animals adopted by the Society for Neuroscience and the Association for Research in Vision and Ophthalmology (Rockville, MD, USA).

### Generation of CRISPR lentiviral particles and subretinal injection

VSV-G pseudotyped lentiviral particles were generated as described(Bainbridge et al., 2001). Subretinal injections were performed under direct retinoscopy thorough an operating microscope. The tip of a 1.5-cm, 34-gauge hypodermic needle (Hamilton) was inserted tangentially through the sclera of the mouse eye, causing a self-sealing wound tunnel. The needle tip was brought into focus between the retina and retinal pigment. Animals received double injections of 2µl each to produce bullous retinal detachments in the superior and inferior hemisphere around the injection sites. Eyes were assigned as treated and (contralateral) control eyes using a randomization software.

### Mammalian antibodies and immunological methods

Fixation and processing cells and mouse tissue sections was as previously described (Zihni et al., 2017b). The following antibodies were used: RPE65 mouse monoclonal (Millipore) 1/300 for immuno-fluorescence; p-MLC S19, mouse monoclonal (Cell Signaling Technology) immunofluorescence 1/100 and immunoblotting 1/1000; Dbl(3), rabbit polyclonal (Santa Cruz Biotechnology) immunofluorescence 1/200; MRCKβ, rabbit polyclonal (Santa Cruz Biotechnology) immunofluorescence 1/200 and immunoblotting 1/500; anti–Cdc42-GTP mouse monoclonal (NewEast Biosciences) immunofluorescence 1/50; N-WASP, rabbit polyclonal (Santa Cruz Biotechnology) immunofluorescence and immunoblotting 1/1000; MerTK rabbit monoclonal (Abcam 52968; recognizes an extracellular N-terminal epitope) immunofluorescence 1/100; IntegrinαVβ5, mouse monoclonal (Clone P1F6) (Abcam 24694) immunofluorescence 1/100; pFAK Y397, rabbit polyclonal (Invitrogen) immunofluorescence 1/100; pSrc Y416, rabbit polyclonal (Cell Signaling Technology) immunofluorescence 1/100; Phalloidin-Atto 647 reagent was obtained from Sigma and diluted 1/1000. Affinity-purified and cross-adsorbed Alexa488-, Cy3- and Cy5-labelled donkey anti-mouse, rabbit, or goat secondary antibodies were from Jackson ImmunoResearch Laboratories (1/300 diluted from 50% glycerol stocks). Affinity-purified HRP-conjugated goat anti mouse and rabbit, and donkey anti goat secondary antibodies 1/5000 were also from Jackson ImmunoResearch Laboratories (1/5000 diluted from 50% glycerol stocks). For immunofluorescence analysis cells and tissues were mounted using Prolong Gold antifade reagent (Life technologies) and imaging was performed using Zeiss 700 and 710 confocal microscopes and a 64x oil lens/NA1.4. Images were processed using Zeiss Zen 2009 and Adobe Photoshop CS5 and 10 software. Co-immunoprecipitation and immunoblotting were carried out using methods previously described and were repeated at least three times (Zihni et al., 2014; Zihni et al., 2017b).

### Isolation of RPE and photoreceptor outer segments

Pig’s eyes were obtained from a slaughterhouse. Briefly in chilled and sterile conditions, each eye was cut into two halves (cornea, including 5mm into the sclera). After removing the lens the eyeball was filled with PBS and neural retina removed using forceps. The PBS was then replaced with 10x Trypsin and incubated at 37oC for 30mins. The trypsin was pipetted several times to dislodge the RPE cells and centrifuged at r.t. 800rpm for 5mins and washed with PBS. The cells were plated into 6-well tissue culture plates with DMEM containing 10% FBS and antibiotics. To isolate POS pig’s eyes were treated as for RPE isolation except that following removal of the lens neural retina was scraped off the tapetum surface and collected in tubes containing homogenization solution. The solution was vigorously shaken for 2min and passed through a double layer gauze to remove tissue fragments. The crude retina preparation was poured into a 30ml ultracentrifuge tube containing chilled continuous sucrose gradients (25-60%) and centrifuged in a Beckman SW-27/SW-32-Ti swing rotor at 25,000rpm at 4oC for 1hour. The appropriate layer was collected and washed three times with centrifugation at 5,000rpm for 10 min each time. The samples were re-suspended in DMEM and stored at -80oC. Isolated POS was labeled before use by centrifuging samples at 6,300 rpm at r.t. for 10min and washing before sonicating for 10 min. FITC label was added to the POS an incubated for 2h whilst rotating 4°C. The samples were then washed 5x in PBS and suspended in appropriate medium.

### Electron Microscopy

All steps were performed at room temperature as previously described (Zihni et al., 2014; Zihni et al., 2017b). Briefly, cell monolayers were fixed in a mixture of 3% (vol/vol) glutaraldehyde and 1% (wt/vol) PFA in 0.08 M sodium cacodylate buffer (CB), pH 7.4, for 2 h at room temperature and left overnight at 4°C. Before osmication, the primary fixative solution was replaced by a 0.08 M cacodylate buffered solution of 2.5% glutaraldehyde and 0.5% (wt/vol) tannic acid. After two brief rinses in CB, specimens were osmicated for 2 h in 1% (wt/vol) aqueous osmium tetroxide, dehydrated by 10-min incubations in 50%, 70%, 90%, and three times 100% ethanol. Semithin sections (0.75μm) for light microscopy and ultrathin sections (50–70 nm) for electron microscopy were cut from sawed out blocks with diamond knives (Diatom; Leica). Semithin sections were stained with 1% toluidine blue/borax mixture at 60°C and ultrathin sections were stained with Reynold’s lead citrate. Stained ultrathin sections were examined in a transmission electron microscopy (1010; JEOL) operating at 80 kV and images were recorded using an Orius B digital camera and Digital Micrograph (Gatan, Inc.)

### Statistical Analysis

Photoreceptor outer segments (POS) internalized into the cell body was measured in Zeiss 700 confocal Z-sections by analyzing FITC-labelled POS particles using Zeiss Zen2000 imaging software or by analyzing Gold-labelled POS by transmission electron microscopy. Percentage of cells with attached POS-FITC was quantified using confocal xy section ranging from the apical cortex to the basal cortex (using phalloidinCy5 as a cortex marker). Cells that contained either externally attached POS-FITC and/or internally attached POS-FITC at >=1 were scored at positive. For in vivo analysis Microvillar organization was measured in electron micrographs using Image-J software. Sheets of microvilli comprising 0, 0-5 or greater than 5, width of microvilli at nm intervals were measured as units every 1.5μm image width. Internalized phagosomes and photoreceptor outer segment fragmentation were measured in electron micrographs using Zeiss Zen2000 software in approximately 20μm fields. External POS fragments were quantified as the total number in contact with the RPE apical membrane/20μm as a sample value of the total neural retinal. ONL thickness, OLM intensity and ratio of apical:basal F-actin cortex intensity were measure using confocal immunofluorescence sections at approximately 200μm widths of retinal tissue. Immunostaining pixel intensity was measured using ImageJ software. For each measurement background was measured and subtracted from the sample value. The control data sets for the immuno-fluorescence experiments Figure S1d,e and 7f,g are the same data set as controls in Figure 1c,d as Control, MRCKβ 1, MRCKβ 2 and Dbl3 were carried out as part of the same independent sets of experiments). For localization of signaling proteins at nascent protrusions xy immunofluorescence confocal sections were measure at the apical membrane of POS stimulated cells both at areas where POS was absent and attached in the same population as an internal control (CF1). Where nascent membrane protrusions were higher than normal microvilli protrusions the membrane was scanned an appropriate number of equal serial sections. For scatter plots of nascent protrusion αvβ5 integrin, pMLC, vinculin, talin individual data points were generated by subtracting first background then regions of the apical membrane that were not attached to POS from attached regions, in the same population as an added internal control. Therefore, graphical Y-axis 0 values = labelling intensity of apical membrane devoid of POS attachment, as highlighted graphically in Figure S3g. Co-localization coefficients were measured in Zeiss confocal EY-scans using Zeiss Zen 2000 co-localization software that applies the principal of the Pearson’s coefficient to selected subcellular structures and using cross hairs function to subtract background from each channel. Co-localization coefficients at specifically nascent protrusions, total cups or POS-signaling plaque contacts at cups were measured by selecting these areas, indicated by colored outlines. The control data sets for THP-1-derived macrophage experiments in Figure 4n are the same data set as controls in Figure 2i as Control, MRCKβ and Dbl3 siRNAs were carried out as part of the same independent sets of experiments. For the quantifications shown, the provided n values refer to independent experiments and the numbers added in brackets in the graphs refer to the total of analyzed cells or number of field widths in electron micrographs and confocal immunofluorescence of mouse retinal tissue. Statistical significance was tested in experiments using two-tailed Student’s t-tests with an n value of at least 3.

## ACKNOWLEDGMENTS

This work was supported by grants from the BBSRC (BB/N014855/1 and BB/N001133/1), Moorfields Eye Charity and the Rosetrees Trust to KM, MSB and CZ; and Retina UK, NIHR Biomedical Research Centre at Moorfields Eye Hospital NHS Foundation Trust support to RRA; and the UCL Institute of Ophthalmology.

---

**Figure S1.**
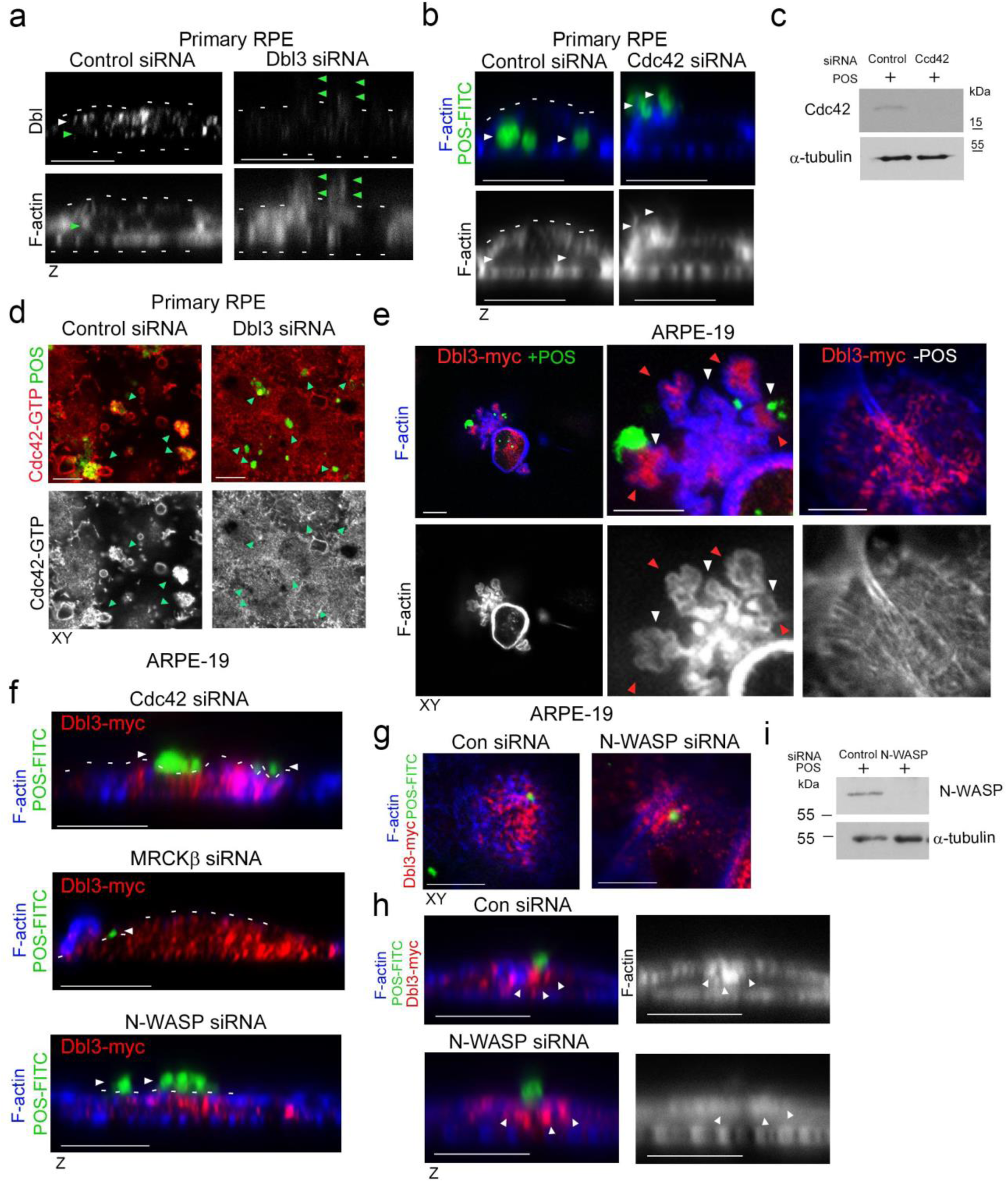
Cdc42 drives protrusion induction and cup maturation. **a,** Confocal z scanning analysis of siRNA-mediated knockdown of Dbl3 in primary porcine RPE cultures treated with POS-FITC for 1h followed by 2h chase. Note, in cells treated with Dbl3 siRNA apical Dbl3 expression is lost. White dashes highlight apical F-actin cortex. Dbl3 knock down also results in stalled f-actin based protrusions highlighted by green arrows. Green arrows in the control siRNA treated, Dbl3 expressing cells highlight the position of internalized POS **b.c,** siRNA-mediated knock-down of Cdc42 in primary porcine RPE cultures treated with POS-FITC for 1h followed by 2h chase inhibits phagocytosis as determined by confocal z-section analysis of porcine primary RPE cells. **d,** Active Cdc42-GTP at nascent protrusions is inhibited by siRNA-mediated knock-down of Dbl3. Note, green arrow heads highlight POS-membrane bound sites. **e,** Confocal xy scanning analysis of ARPE-19 cells expressing Dbl3-myc with or without POS stimulation. Note, POS adhesion stimulates Dbl3-driven actin-rich cup formation but Dbl3-myc exogenous expression only does not stimulate cups. White arrows indicate cups attached to POS. Red arrows highlight Dbl3-myc localization at cup tips. **f,** Confocal z-scanning analysis showing effects of siRNA-mediated knock-down of Cdc42 and its effectors in Dbl3-myc expressing ARPE-19 cells. Note, knockdown of Cdc42 signaling in Dbl3-myc driven phagocytosis result in an inhibition of phagocytosis. **g-i,** Confocal xy and z sections of ARPE-19 cells with N-WASP knockdown expressing Dbl3-myc and stimulated with POS-FITC for 30minutes reveals cells require N-WASP for F-actin polymerization during protrusion induction. White arrows highlight loss of Dbl3-myc induced F-actin staining revealed by z-section analysis at sites of POS attachment following N-WASP siRNA-mediated knock-down. All analyzes are based on n=3 independent experiments (except for Cdc42 siRNA experiments shown in b where n=4).

**Figure S2.**
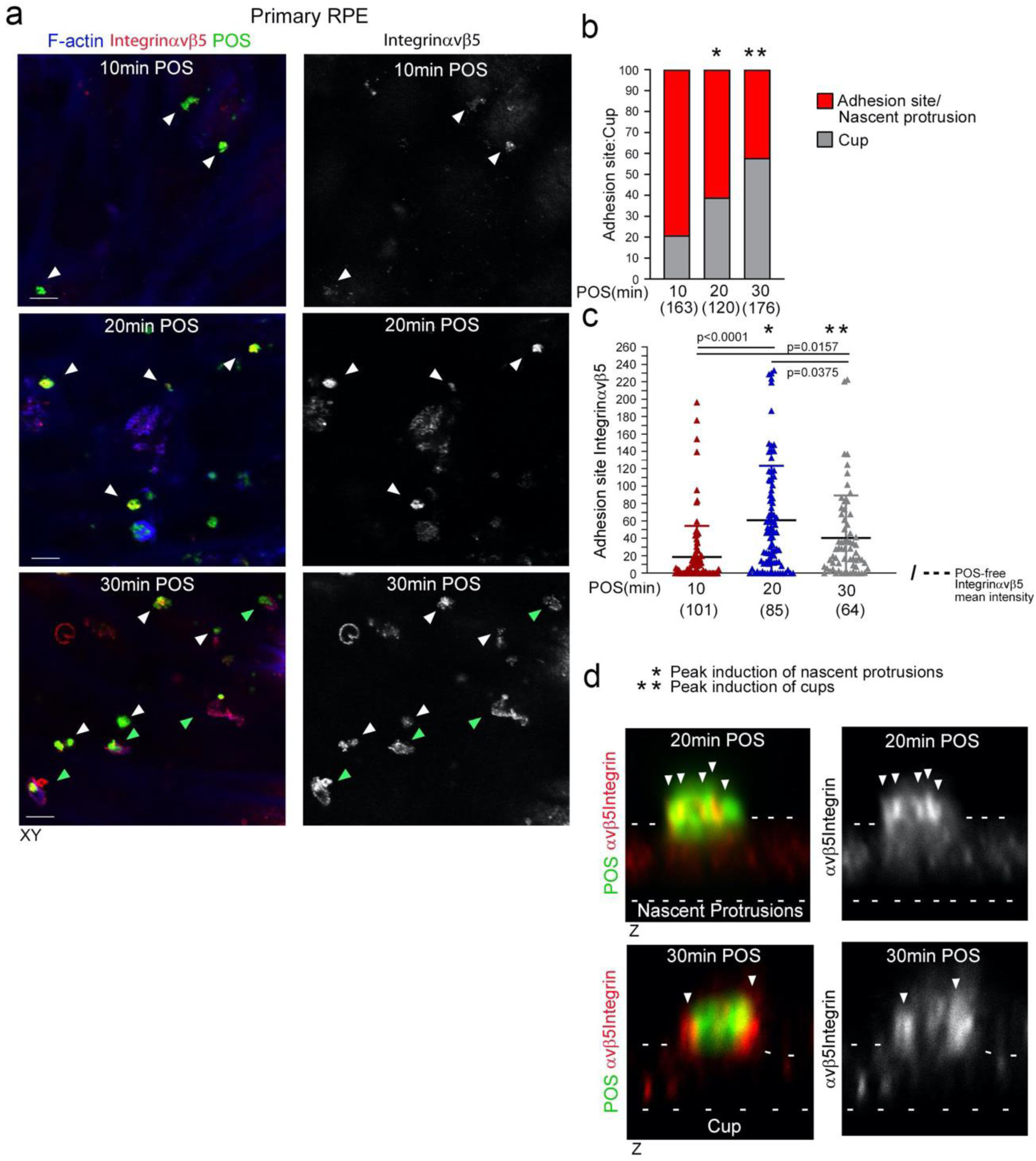
POS-membrane contacts induce membrane deformation. **a-c**, Time course of POS incubation with primary RPE cells reveals that adhesion sites are enriched with receptor within 10min of POS addition, as determined by adhesion receptor αvβ5 integrin staining and peak with respect to mean labelling intensity at 20min. **d,** At 20min membrane deformation results in protrusions that transform into cups that wrap around POS at 30min. Note, green arrows highlight cups. Protein staining was quantified as mean intensity. Scale bars represent 10μm. All analyzes are based on n=3 independent experiments. and show the data points, means ±1SD, the total number of cells analyzed for each type of sample across all experiments, and p-values derived from t-tests.

**Figure S3.**
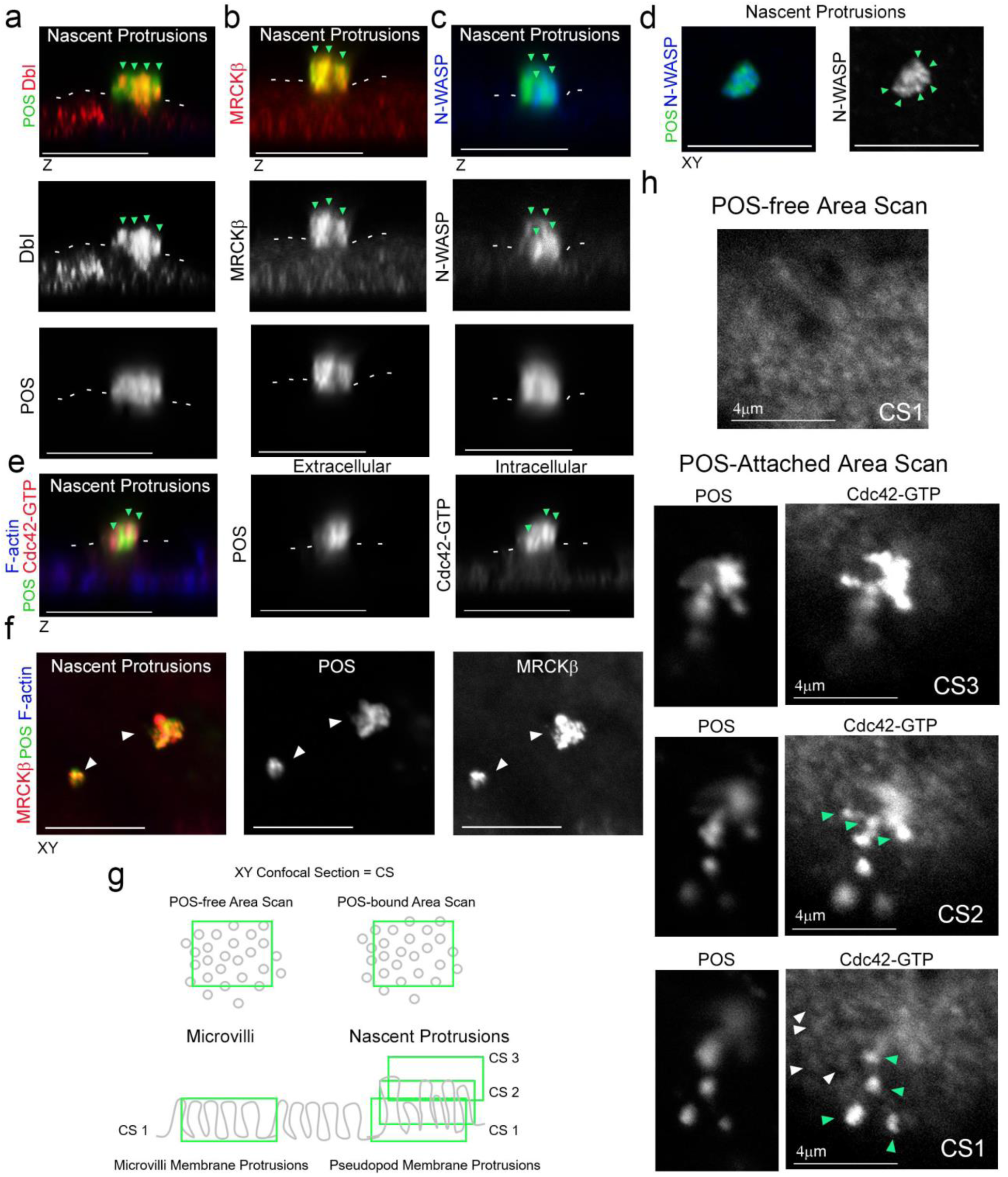
RPE nascent protrusions are rich in Cdc42-signaling machinery. **a-c,e,** Confocal z-section analysis reveals that POS binding to the apical membrane of RPE, induces membrane protrusions rich in Dbl3, Cdc42-GTP, MRCKβ and N-WASP. White dashes highlight the normal apical membrane. **d,** Confocal xy section of N-WASP localization at nascent protrusions. **f,** Confocal xy analysis of POS-induced nascent protrusions reveals that the size of induced protrusions correlates with the size of POS particle. **g,** Schematic diagram illustrating method of quantifying mean labelling intensity of protrusion proteins using Cdc42-GTP as an example. Briefly, confocal xy sections were scanned at the apical membrane surface of RPE cells (Confocal Scan 1, CS1) both unattached and POS attached. Since protrusions are at an increased hight compared to normal apical brush border microvilli further serial scans were taken as appropriate to cover the protrusion length and values were averaged. **h,** At CS1 Cdc42-GTP enrichment is observed at single protrusions (green arrows) at a similar diameter to surrounding non-POS attached brush border structures (white arrows).

**Figure S4.**
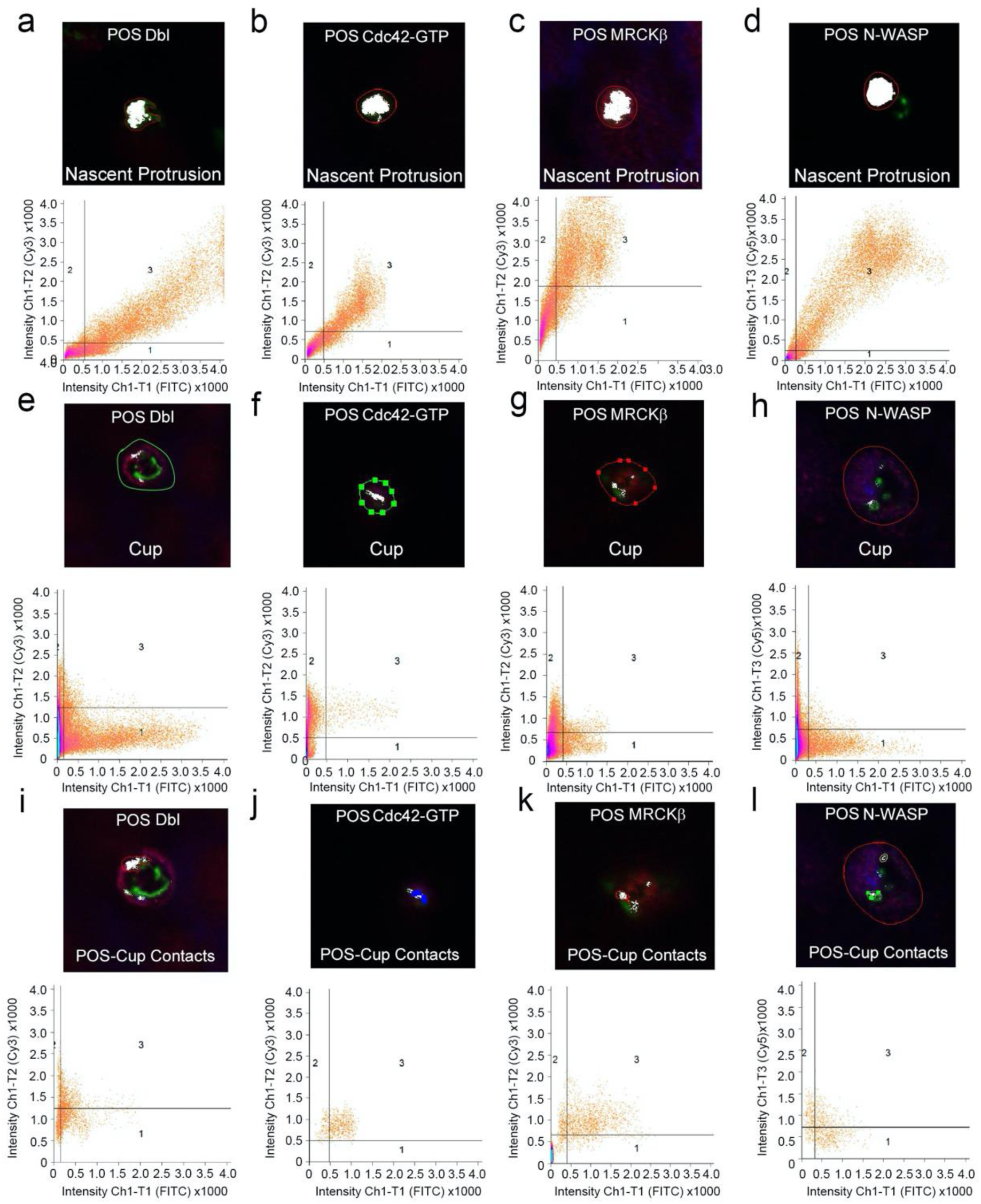
POS co-localization with Cdc42 network at protrusions and Cups. **a-d**, Primary porcine RPE cells were treated with POS-FITC for 20min or **e-l,** 30min and analyzed by confocal microscopy using co-localization software based on the Pearson’s correlation coefficient. Co-localization was calculated in grid 3 of the scatter charts. White represents co-localization of POS-FITC with either Dbl, Cdc42-GTP, MRCKβ or N-WASP, which is nearly total at nascent protrusions and only partial, at the particle contact interface in the cup configuration. Colored outlines of nascent protrusions, cups or POS-Cup contacts represent calculation areas of co-localization coefficients. All analyzes are based on n=3 independent experiments.

**Figure S5.**
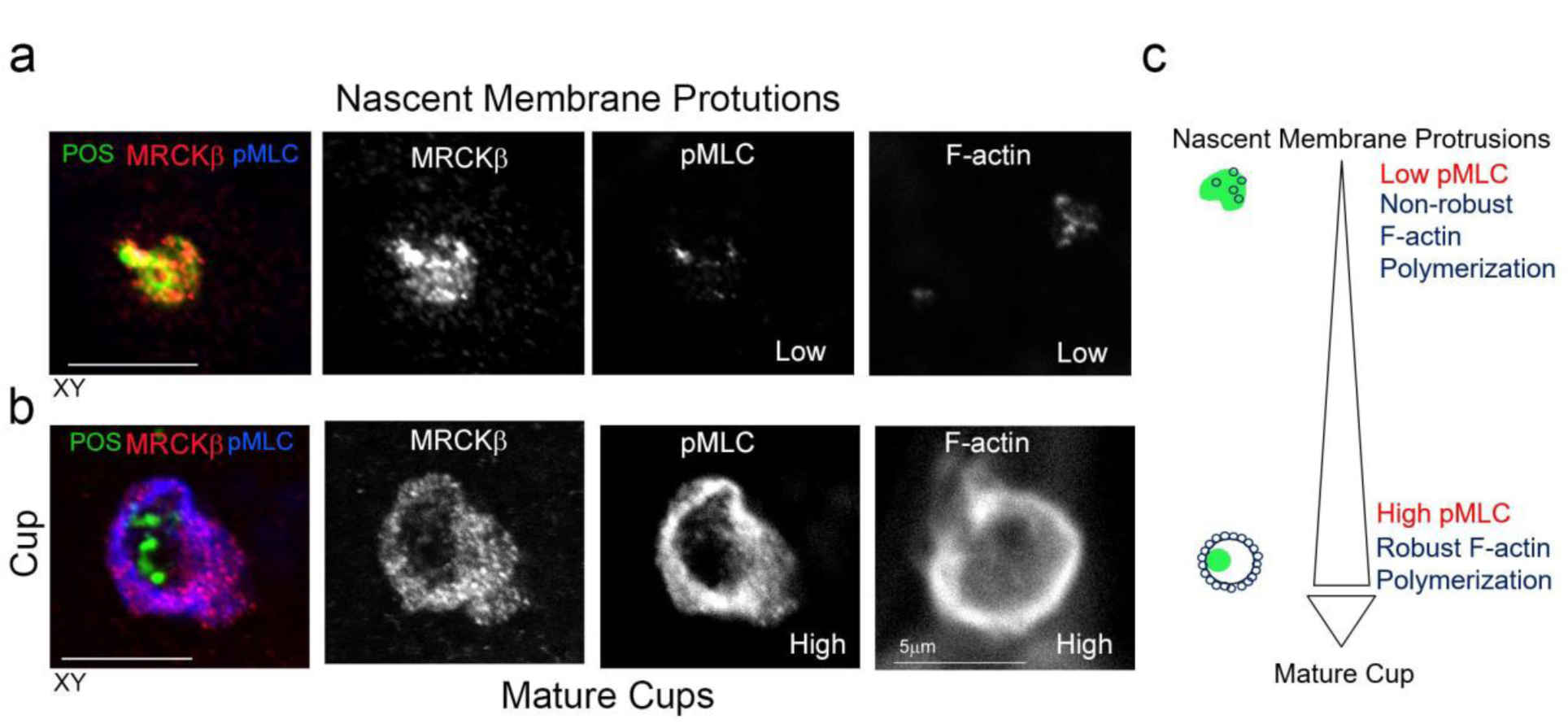
pMLC levels increases during protrusion maturation to cups. **a-c,** Confocal xy sections of primary porcine RPE cells treated with POS-FITC for up to 30min reveal that nascent protrusions display low pMLC activity and weak F-actin staining that become significantly more intense in mature cups. Note, the increase in myosin II activity during protrusion to cup maturation occurs concomitantly with robust actin polymerization.

**Figure S6.**
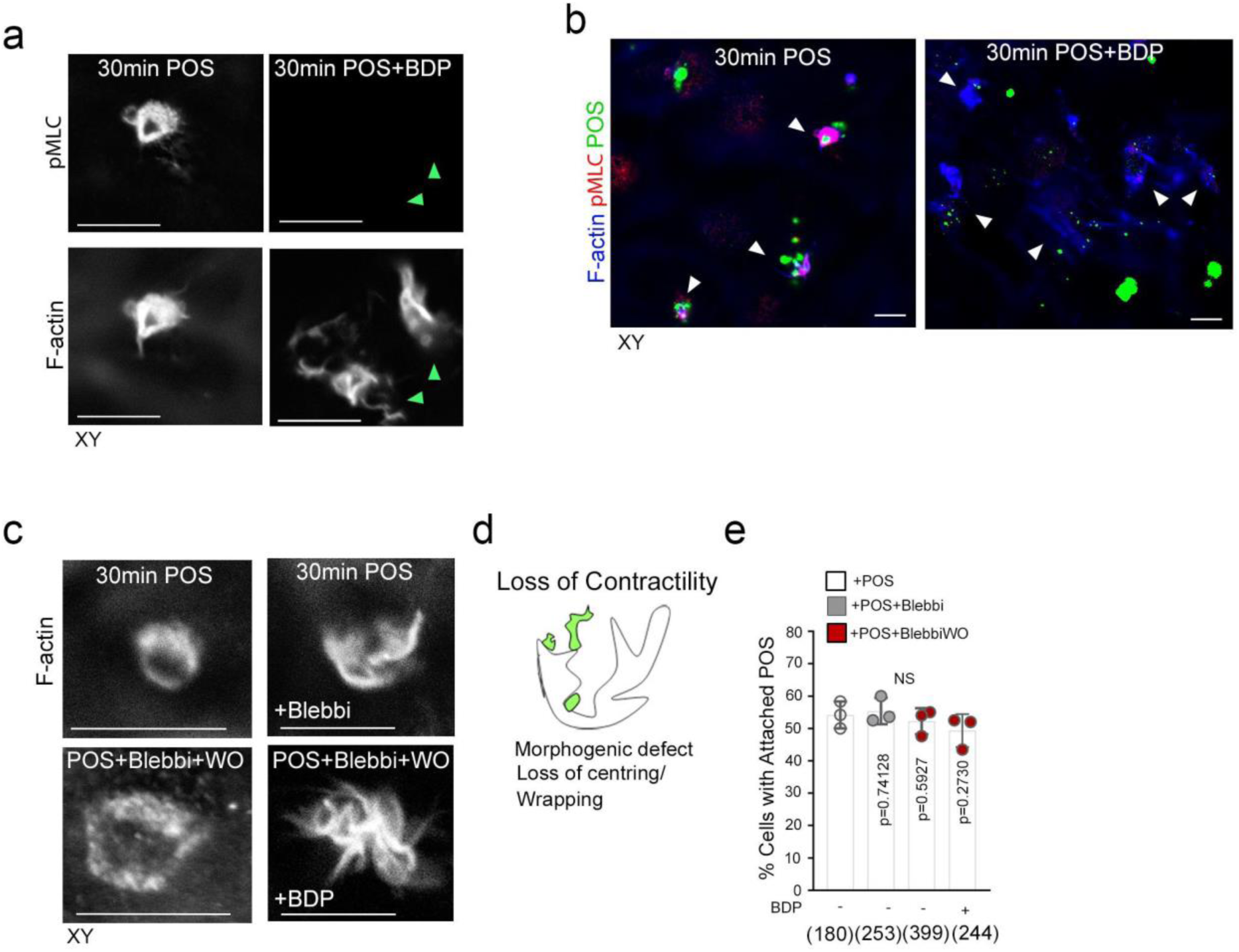
Cup wrapping around POS requires MRCKβ/myosin-II. **a,b** Confocal xy scans of porcine primary RPE cells treated with POS for 30 minutes reveal cups encircling the particles with increased pMLC activity compared to POS-free segments of apical membrane. Treatment of cells with the MRCK inhibitor BDP5290 results in pseudopods negative for pMLC that are morphogenetically defective and unable to wrap around POS. Note, overall efficiency of POS binding to RPE cells is unaltered. **c-e,** Confocal xy scans of primary cells treated with POS and blebbistatin reveal pseudopods with a morphological defect having lost their cylindrical cup-like confirmation similar to MRCK inhibition that is rapidly reversed after 2h blebbistatin washout, only in the presence of active MRCKβ. Note, treatment of cells with blebbistatin or BDP5290 does not affect binding efficiency of POS.

**Figure S7.**
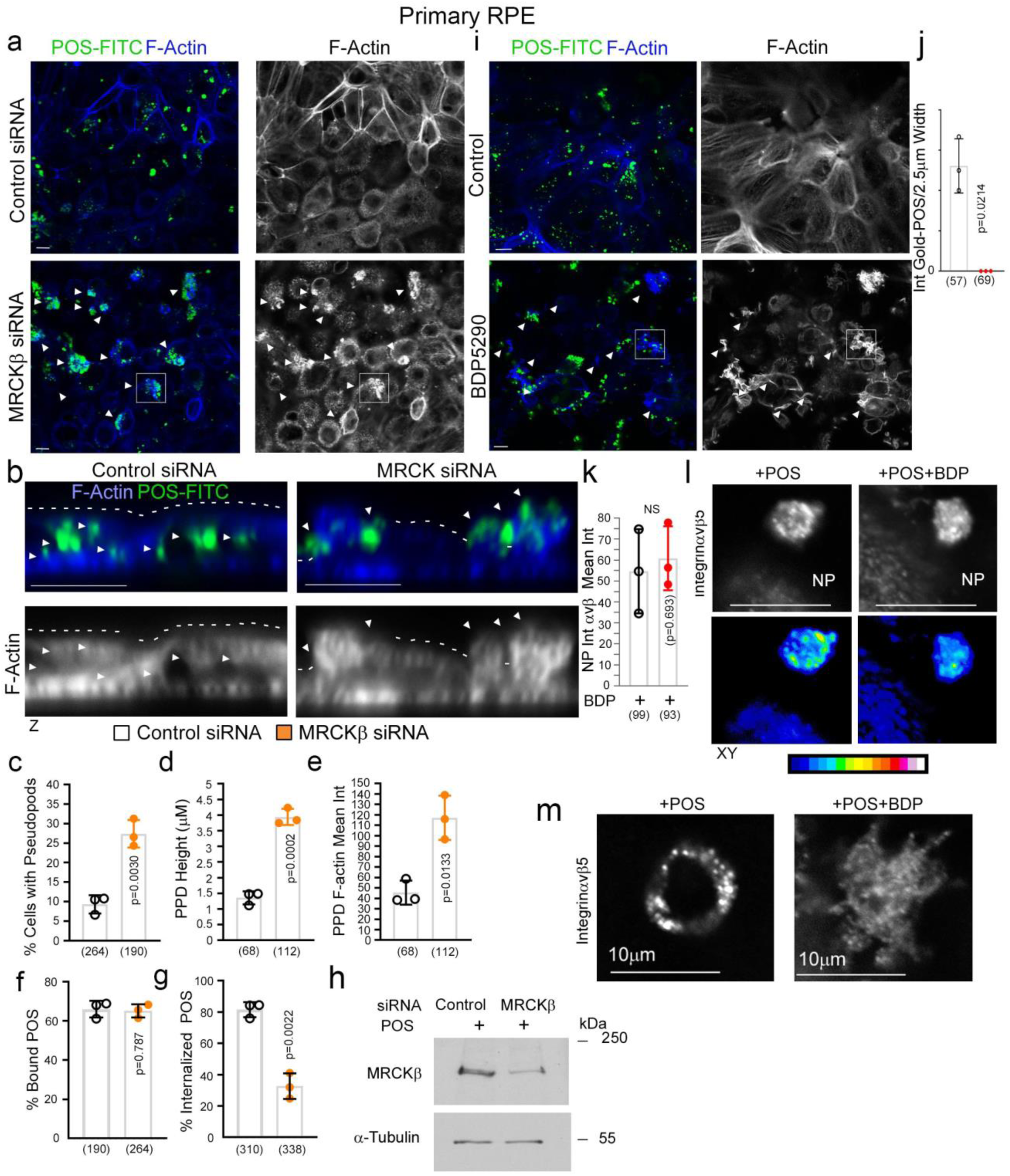
MRCKβ regulates actin dynamics during cup maturation. **a-h,** siRNA-mediated knockdown of MRCKβ in primary porcine RPE cells leads to prolonged, longer and disordered pseudopods with increased F-actin labelling. **i,** Inhibition of MRCKβ kinase activity using BDP5290 results in a membrane remodeling defect in a similar manner to knock-down of MRCKβ. Note, in both treatments total adhesion of POS to RPE cells is unaffected although POS is often misaligned and not centered at extended pseudopods in either MRCKβ siRNA-treated or BDP5290 treated cells as indicated by white arrows in the confocal xy-scanning sections. **j,** inhibition of MRCKβ kinase activity inhibits phagocytosis as determined by immuno-Gold POS labelled TEM analysis. **k, l,** Confocal xy analysis of primary RPE nascent protrusions with or without MRCKβ inhibition. Upper panels are grey scale images of integrin αvβ5 localization as used for measurements of mean intensity values, and lower panels are resulting heatmaps of the integrin αvβ5 staining to highlight enrichment along protrusions. **m,** Confocal xy analysis of integrin αvβ5 distribution in cups with or without MRCKβ inhibition. Note, inhibition of MRCKβ-driven actomyosin contractility results in loss of integrin αvβ5 clustering along the deformed pseudopods that do not mature into normal cylindrical cups as highlighted by F-actin staining. Scale bars represent 10μm. All quantifications are based on n=3 independent experiments and show the data points, means ±1SD, the total number of cells analyzed for each type of sample across all experiments, and p-values derived from t-tests.

**Figure S8.**
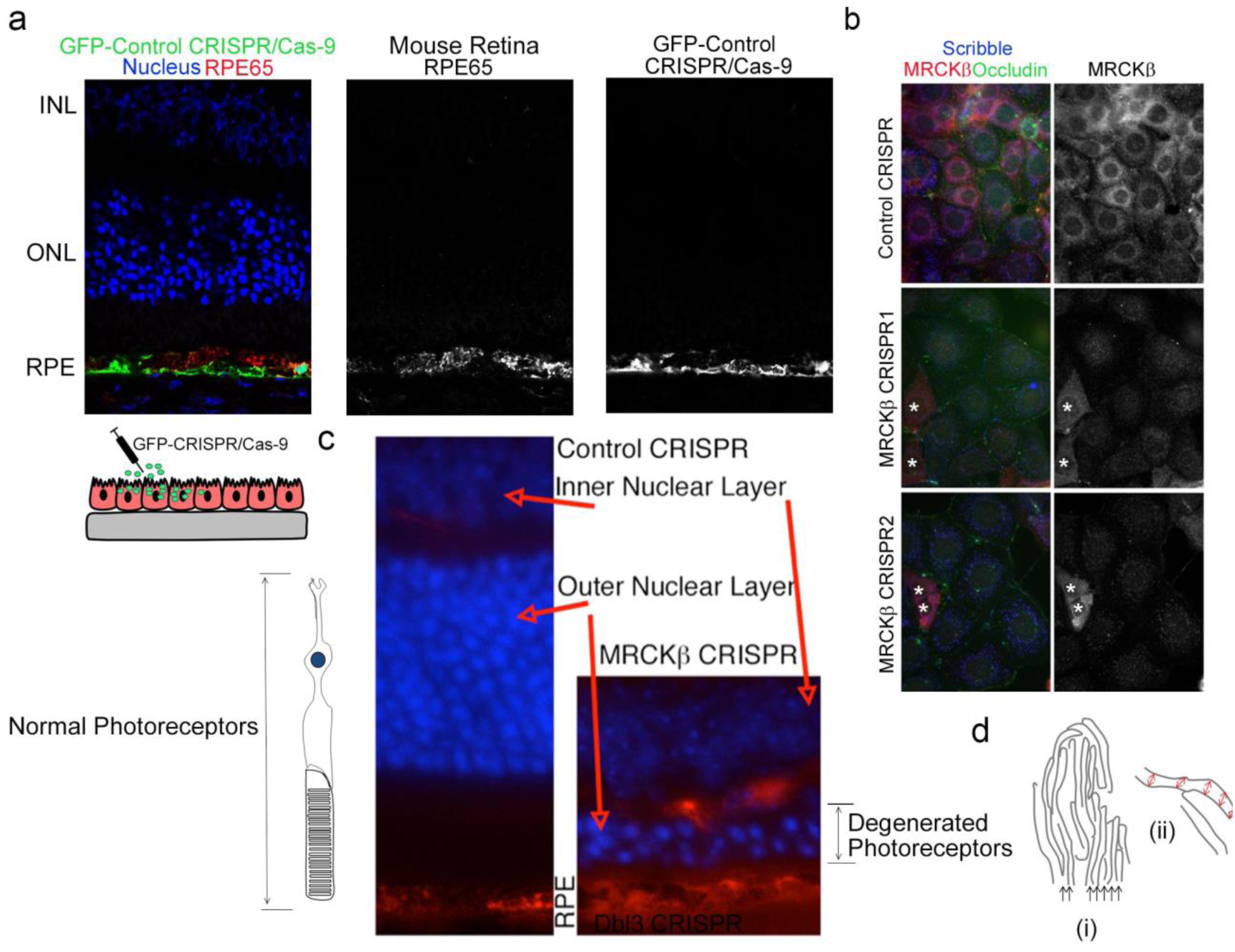
MRCKβ is required for retinal integrity and function in vivo. **a**, Subretinally injected lentiviral vectors specifically infect the RPE. Shown is GFP expression upon transduction with a control vector, combined with staining for the RPE marker RPE65 and DNA. **b**, MRCKβ expression upon transduction with control CRISPR or two distinct viruses targeting the MRCKβ gene. White asterisk signify cells still expressing MRCKβ that were included as controls for image exposure. **c**, Frozen sections of mouse eyes 21 days after injection with either control or MRCKβ-targeting viruses were stained for RPE65 and DNA. Note, MRCKβ knockout leads to a pronounced retinal degeneration as determined by extensive thinning of the ONL. **d**, Schematic diagram illustrating measurement parameters of RPE apical microvilli to determine morphology and order. Red arrows represent measurements of width of each MV at regular intervals and black arrows highlight individual microvilli strands. **e-g,** Confocal immunofluorescence analysis of retinal sections from mice injected with either control, or MRCKβ 2, knockout vector that targets a distinct region of the MRCKβ gene reveal, as with MRCKβ 1 knock out vector, no significant difference in the overall retinal structure after 7 days but reduced apical F-actin staining, with reference to basal F-actin, in the MRCKβ-deficient RPE. White arrows highlight apical F-actin cortex. Protein staining was quantified as mean intensity. Quantifications show means±1SD, n=3, p-values derived from t-tests.

**Figure S9.**
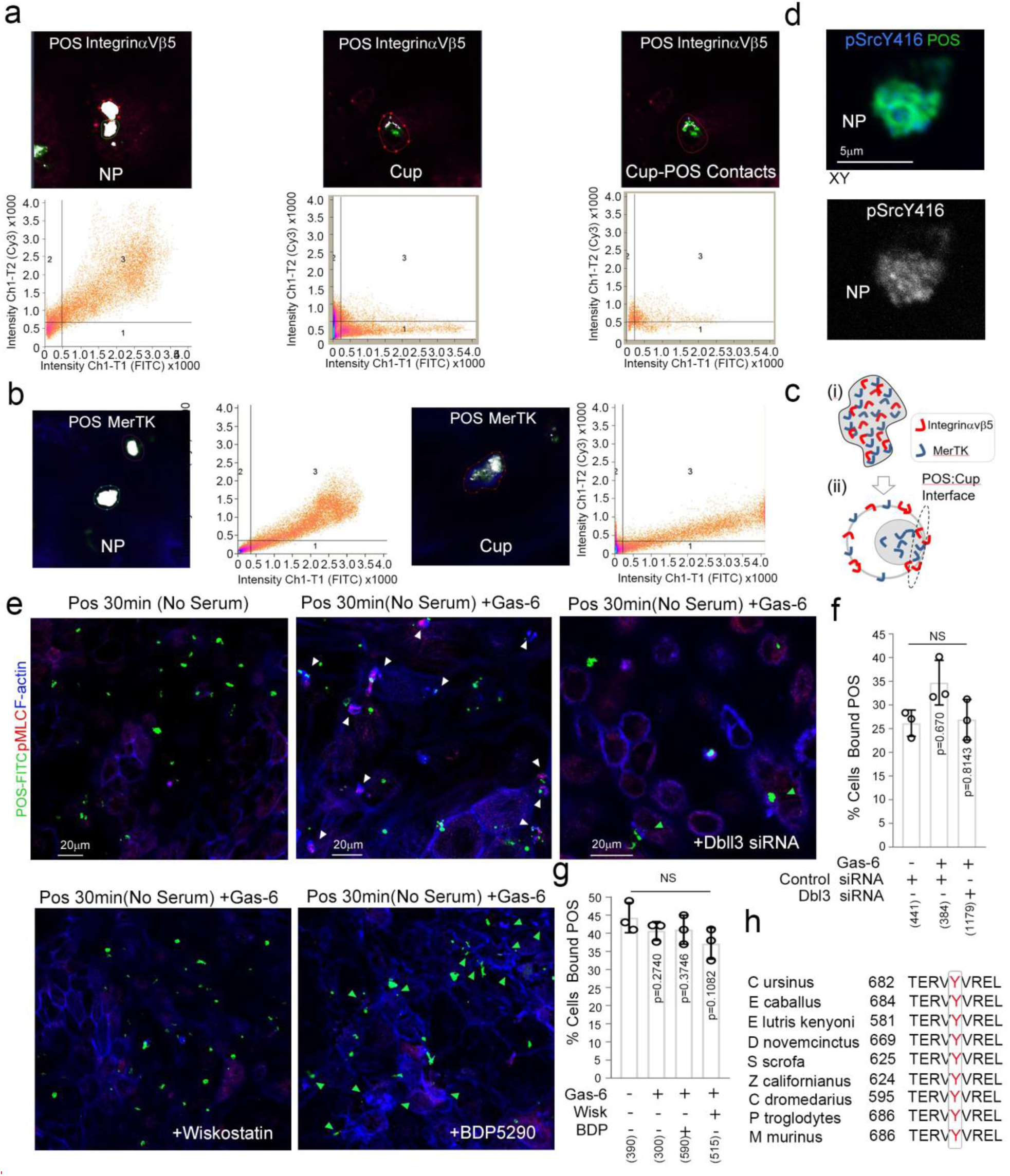
MerTK-signaling during particle wrapping. **a-c,** Primary porcine RPE cells were analyzed by confocal xy scanning microscopy for localization of αvβ5 and MerTK receptors relative to POS using colocalization software based on the Pearson’s correlation coefficient. Grid 3 of the scatter charts contains the colocalization data calculations. Colocalization of POS-FITC with either integrin αvβ5, or MerTK is depicted in white. Note, colocalization of POS receptors with membrane-bound POS is nearly total at nascent protrusions but with integrin αvβ5 only partial at cups where co-localization occurs at POS contact sites at the internal side of cups. MerTK also localizes at cups and contact sites but more substantially with total POS. Colored outlines of nascent protrusions, cups or POS-cup contacts represent areas used for calculating of colocalization coefficients. **d,** Confocal microscopy analysis of primary RPE treated with POS-FITC revealed activated MerTK receptor-associated tyrosine kinase Src (pSrcY416) at nascent protrusions. All analyzes are based on n=3 independent experiments. **e-g,** Confocal xy analysis of cup morphogenesis, pMLC staining and POS centering in primary RPE treated with POS-Gas-6 with or without the N-WASP inhibitor Wiskostatin, MRCK inhibitor BDP5290 or siRNA-mediated Dbl3 knockdown. Note, Gas-6 with POS drives cup maturation and wrapping around POS and increased pMLC. White arrows highlight cups with centered POS and green arrows highlight protrusions with misaligned POS. Treatment of cells with Gas-6 or siRNAs did not significantly affect efficiency of POS-FITC binding to the cell membrane. **h,** Sequence alignment of the Dbl GEF motif for activation by c-Src tyrosine phosphorylation in diverse mammalian species demonstrating high evolutionary conservation. All quantifications are based on n=3 independent experiments and show the data points, means ±1SD, the total number of cells analyzed for each type of sample across all experiments, and p-values derived from t-tests.

**Figure S10.**
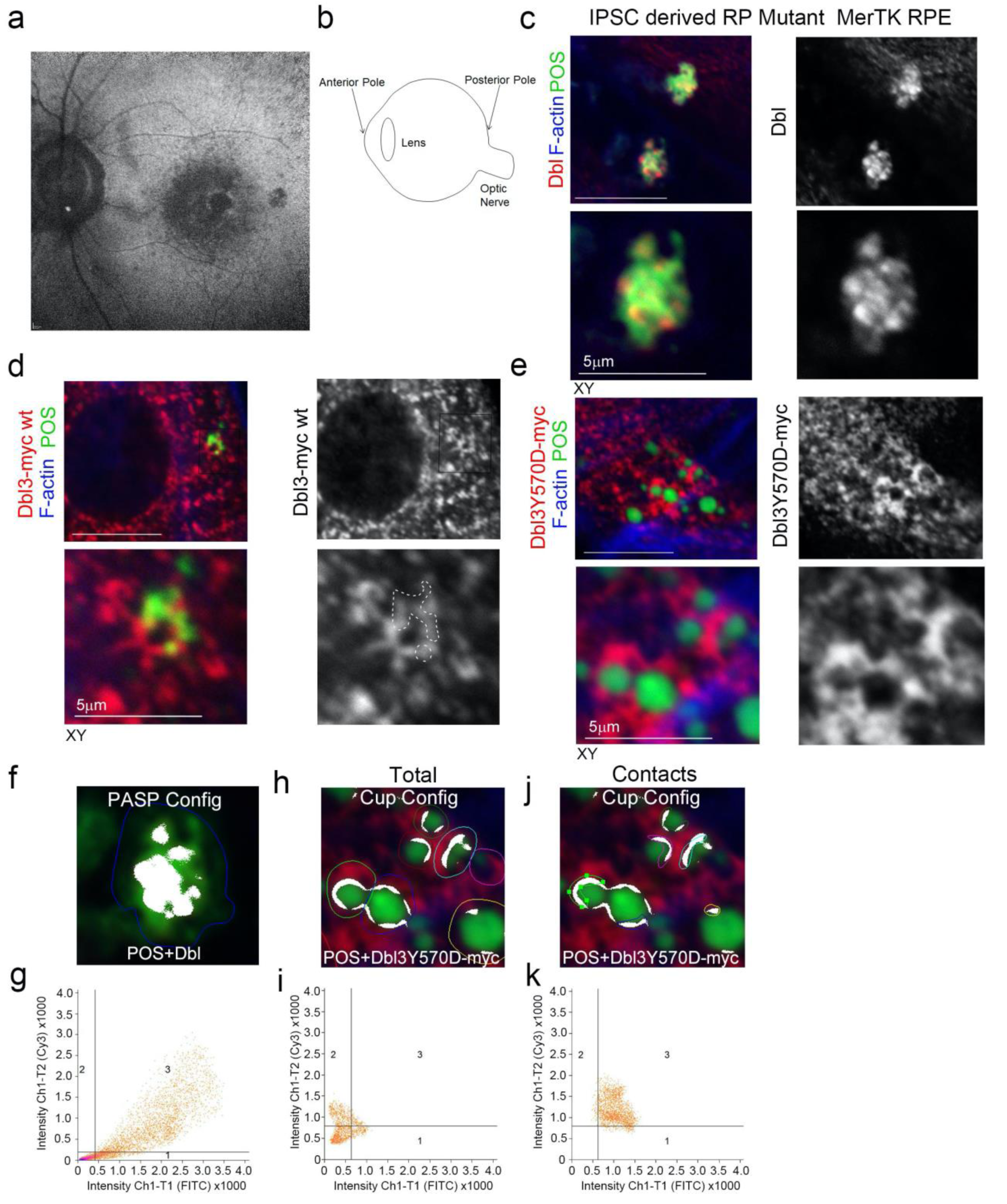
Dbl3Y570D rescues phagocytosis in MerTK-deficient RPE. **a,b,** Autofluorescence image of a magnified area of the posterior pole. The speckled darker area at the central macula represents a dropout of RPE cells from an individual carrying a loss of function MerTK mutation that causes retinitis pigmentosa (RP) and vision loss. **c,** confocal xy scanning fluorescence image of RPE cells differentiated from iPSCs derived from an RP individual reveals an accumulation of adhesion sites with enriched Dbl3 at POS contact sites but no wrapping of POS. **d**, Expression of wild type Dbl3-myc does not promote significant maturation of adhesion sites to cups. **e,** Expression of Dbl3Y570D-myc tyrosine phospho-mimetic mutant promotes efficient nascent adhesion to cup maturation despite loss of upstream MerTK function. **f-k,** RP individual iPSC-derived RPE expressing different forms of Dbl analyzed by confocal xy scanning microscopy using co-localization software based on the Pearson’s correlation coefficient. Note, coloured outlines in the confocal xy-section images represent areas of calculation. Dbl3Y570D fully restores progression from complete colocalization at adhesion sites to POS-membrane Dbl3 interface contacts in the cup configuration. Note, grid 3 of the scatter graph represents colocalization data calculations and Dbl co-localization with POS is highlighted in white. All quantifications are based on n=3 independent experiments and show the data points, means ±1SD, the total number of cells analyzed for each type of sample across all experiments, and p-values derived from t-tests.

## REFERENCES

1. Ablonczy, Z., M. Dahrouj, P.H. Tang, Y. Liu, K. Sambamurti, A.D. Marmorstein, and C.E. Crosson. 2011. Human retinal pigment epithelium cells as functional models for the RPE in vivo. Invest Ophthalmol Vis Sci. 52:8614–8620.

2. Ardura, J.A., G. Rackov, E. Izquierdo, V. Alonso, A.R. Gortazar, and M.M. Escribese. 2019. Targeting Macrophages: Friends or Foes in Disease? Front Pharmacol. 10:1255.

3. Bainbridge, J.W., C. Stephens, K. Parsley, C. Demaison, A. Halfyard, A.J. Thrasher, and R.R. Ali. 2001. In vivo gene transfer to the mouse eye using an HIV-based lentiviral vector; efficient long-term transduction of corneal endothelium and retinal pigment epithelium. Gene Ther. 8:1665–1668.

4. Barger, S.R., N.C. Gauthier, and M. Krendel. 2020. Squeezing in a Meal: Myosin Functions in Phagocytosis. Trends Cell Biol. 30:157–167.

5. Bonilha, V.L., M.E. Rayborn, S.K. Bhattacharya, X. Gu, J.S. Crabb, J.W. Crabb, and J.G. Hollyfield. 2006. The retinal pigment epithelium apical microvilli and retinal function. Adv Exp Med Biol. 572:519–524.

6. Brosius Lutz, A., W.S. Chung, S.A. Sloan, G.A. Carson, L. Zhou, E. Lovelett, S. Posada, J.B. Zuchero, and B.A. Barres. 2017. Schwann cells use TAM receptor-mediated phagocytosis in addition to autophagy to clear myelin in a mouse model of nerve injury. Proc Natl Acad Sci U S A. 114:E8072–E8080.

7. Caron, E., and A. Hall. 1998. Identification of two distinct mechanisms of phagocytosis controlled by different Rho GTPases. Science. 282:1717–1721.

8. Carvalho, K., J. Lemiere, F. Faqir, J. Manzi, L. Blanchoin, J. Plastino, T. Betz, and C. Sykes. 2013. Actin polymerization or myosin contraction: two ways to build up cortical tension for symmetry breaking. Philos Trans R Soc Lond B Biol Sci. 368:20130005.

9. Case, L.B., and C.M. Waterman. 2015. Integration of actin dynamics and cell adhesion by a three-dimensional, mechanosensitive molecular clutch. Nat Cell Biol. 17:955–963.

10. Chaitin, M.H., and M.O. Hall. 1983. Defective ingestion of rod outer segments by cultured dystrophic rat pigment epithelial cells. Invest Ophthalmol Vis Sci. 24:812–820.

11. Chen, C.S. 2008. Mechanotransduction - a field pulling together? J Cell Sci. 121:3285–3292.

12. Clarke, M., U. Engel, J. Giorgione, A. Muller-Taubenberger, J. Prassler, D. Veltman, and G. Gerisch. 2010. Curvature recognition and force generation in phagocytosis. BMC Biol. 8:154.

13. D’Cruz, P.M., D. Yasumura, J. Weir, M.T. Matthes, H. Abderrahim, M.M. LaVail, and D. Vollrath. 2000. Mutation of the receptor tyrosine kinase gene Mertk in the retinal dystrophic RCS rat. Hum Mol Genet. 9:645–651.

14. del Rio, A., R. Perez-Jimenez, R. Liu, P. Roca-Cusachs, J.M. Fernandez, and M.P. Sheetz. 2009. Stretching single talin rod molecules activates vinculin binding. Science. 323:638–641.

15. Edwards, R.B., and R.B. Szamier. 1977. Defective phagocytosis of isolated rod outer segments by RCS rat retinal pigment epithelium in culture. Science. 197:1001–1003.

16. Etienne-Manneville, S., and A. Hall. 2002. Rho GTPases in cell biology. Nature. 420:629–635.

17. Finnemann, S.C. 2003. Focal adhesion kinase signaling promotes phagocytosis of integrin-bound photoreceptors. EMBO J. 22:4143–4154.

18. Finnemann, S.C., V.L. Bonilha, A.D. Marmorstein, and E. Rodriguez-Boulan. 1997. Phagocytosis of rod outer segments by retinal pigment epithelial cells requires alpha(v)beta5 integrin for binding but not for internalization. Proc Natl Acad Sci U S A. 94:12932–12937.

19. Finnemann, S.C., and E. Rodriguez-Boulan. 1999. Macrophage and retinal pigment epithelium phagocytosis: apoptotic cells and photoreceptors compete for alphavbeta3 and alphavbeta5 integrins, and protein kinase C regulates alphavbeta5 binding and cytoskeletal linkage. J Exp Med. 190:861–874.

20. Freeman, S.A., and S. Grinstein. 2014. Phagocytosis: receptors, signal integration, and the cytoskeleton. Immunol Rev. 262:193–215.

21. Georgiadis, A., M. Tschernutter, J.W. Bainbridge, K.S. Balaggan, F. Mowat, E.L. West, P.M. Munro, A.J. Thrasher, K. Matter, M.S. Balda, and R.R. Ali. 2010. The tight junction associated signalling proteins ZO-1 and ZONAB regulate retinal pigment epithelium homeostasis in mice. PLoS One. 5:e15730.

22. Grinnell, F. 1984. Fibroblast spreading and phagocytosis: similar cell responses to different-sized substrata. J Cell Physiol. 119:58–64.

23. Gupta, M., X. Qi, V. Thakur, and D. Manor. 2014. Tyrosine phosphorylation of Dbl regulates GTPase signaling. J Biol Chem. 289:17195–17202.

24. Hall, M.O., M.S. Obin, A.L. Prieto, B.L. Burgess, and T.A. Abrams. 2002. Gas6 binding to photoreceptor outer segments requires gamma-carboxyglutamic acid (Gla) and Ca(2+) and is required for OS phagocytosis by RPE cells in vitro. Exp Eye Res. 75:391–400.

25. Haukedal, H., and K. Freude. 2019. Implications of Microglia in Amyotrophic Lateral Sclerosis and Frontotemporal Dementia. J Mol Biol. 431:1818–1829.

26. Hoppe, A.D., and J.A. Swanson. 2004. Cdc42, Rac1, and Rac2 display distinct patterns of activation during phagocytosis. Mol Biol Cell. 15:3509–3519.

27. Humphries, J.D., P. Wang, C. Streuli, B. Geiger, M.J. Humphries, and C. Ballestrem. 2007. Vinculin controls focal adhesion formation by direct interactions with talin and actin. J Cell Biol. 179:1043–1057.

28. Inana, G., C. Murat, W. An, X. Yao, I.R. Harris, and J. Cao. 2018. RPE phagocytic function declines in age-related macular degeneration and is rescued by human umbilical tissue derived cells. J Transl Med. 16:63.

29. Jaumouille, V., A.X. Cartagena-Rivera, and C.M. Waterman. 2019. Coupling of beta2 integrins to actin by a mechanosensitive molecular clutch drives complement receptor-mediated phagocytosis. Nat Cell Biol. 21:1357–1369.

30. Jiang, M., J. Esteve-Rudd, V.S. Lopes, T. Diemer, C. Lillo, A. Rump, and D.S. Williams. 2015. Microtubule motors transport phagosomes in the RPE, and lack of KLC1 leads to AMD-like pathogenesis. J Cell Biol. 210:595–611.

31. Kevany, B.M., and K. Palczewski. 2010. Phagocytosis of retinal rod and cone photoreceptors. Physiology (Bethesda). 25:8–15.

32. Kovacs, M., J. Toth, C. Hetenyi, A. Malnasi-Csizmadia, and J.R. Sellers. 2004. Mechanism of blebbistatin inhibition of myosin II. J Biol Chem. 279:35557–35563.

33. Kumfer, K.T., S.J. Cook, J.M. Squirrell, K.W. Eliceiri, N. Peel, K.F. O’Connell, and J.G. White. 2010. CGEF-1 and CHIN-1 regulate CDC-42 activity during asymmetric division in the Caenorhabditis elegans embryo. Mol Biol Cell. 21:266–277.

34. Law, A.L., C. Parinot, J. Chatagnon, B. Gravez, J.A. Sahel, S.S. Bhattacharya, and E.F. Nandrot. 2015. Cleavage of Mer tyrosine kinase (MerTK) from the cell surface contributes to the regulation of retinal phagocytosis. J Biol Chem. 290:4941–4952.

35. Lin, H., and D.O. Clegg. 1998. Integrin alphavbeta5 participates in the binding of photoreceptor rod outer segments during phagocytosis by cultured human retinal pigment epithelium. Invest Ophthalmol Vis Sci. 39:1703–1712.

36. Mao, Y., and S.C. Finnemann. 2012. Essential diurnal Rac1 activation during retinal phagocytosis requires alphavbeta5 integrin but not tyrosine kinases focal adhesion kinase or Mer tyrosine kinase. Mol Biol Cell. 23:1104–1114.

37. Mao, Y., and S.C. Finnemann. 2015. Regulation of phagocytosis by Rho GTPases. Small GTPases. 6:89–99.

38. Marston, D.J., C.D. Higgins, K.A. Peters, T.D. Cupp, D.J. Dickinson, A.M. Pani, R.P. Moore, A.H. Cox, D.P. Kiehart, and B. Goldstein. 2016. MRCK-1 Drives Apical Constriction in C. elegans by Linking Developmental Patterning to Force Generation. Curr Biol. 26:2079–2089.

39. Massol, P., P. Montcourrier, J.C. Guillemot, and P. Chavrier. 1998. Fc receptor-mediated phagocytosis requires CDC42 and Rac1. EMBO J. 17:6219–6229.

40. Miceli, M.V., D.A. Newsome, and D.J. Tate, Jr. 1997. Vitronectin is responsible for serum-stimulated uptake of rod outer segments by cultured retinal pigment epithelial cells. Invest Ophthalmol Vis Sci. 38:1588–1597.

41. Miki, H., T. Sasaki, Y. Takai, and T. Takenawa. 1998. Induction of filopodium formation by a WASP-related actin-depolymerizing protein N-WASP. Nature. 391:93–96.

42. Mullen, R.J., and M.M. LaVail. 1976. Inherited retinal dystrophy: primary defect in pigment epithelium determined with experimental rat chimeras. Science. 192:799–801.

43. Myers, K.V., S.R. Amend, and K.J. Pienta. 2019. Targeting Tyro3, Axl and MerTK (TAM receptors): implications for macrophages in the tumor microenvironment. Mol Cancer. 18:94.

44. Oria, R., T. Wiegand, J. Escribano, A. Elosegui-Artola, J.J. Uriarte, C. Moreno-Pulido, I. Platzman, P. Delcanale, L. Albertazzi, D. Navajas, X. Trepat, J.M. Garcia-Aznar, E.A. Cavalcanti-Adam, and P. Roca-Cusachs. 2017. Force loading explains spatial sensing of ligands by cells. Nature. 552:219–224.

45. Pasapera, A.M., I.C. Schneider, E. Rericha, D.D. Schlaepfer, and C.M. Waterman. 2010. Myosin II activity regulates vinculin recruitment to focal adhesions through FAK-mediated paxillin phosphorylation. J Cell Biol. 188:877–890.

46. Ramsden, C.M., B. Nommiste, R.L. A, A.F. Carr, M.B. Powner, J.K.S. M, L.L. Chen, M.N. Muthiah, A.R. Webster, A.T. Moore, M.E. Cheetham, L. da Cruz, and P.J. Coffey. 2017. Rescue of the MERTK phagocytic defect in a human iPSC disease model using translational read-through inducing drugs. Sci Rep. 7:51.

47. Reymann, A.C., R. Boujemaa-Paterski, J.L. Martiel, C. Guerin, W. Cao, H.F. Chin, E.M. De La Cruz, M. Thery, and L. Blanchoin. 2012. Actin network architecture can determine myosin motor activity. Science. 336:1310–1314.

48. Shelby, S.J., K.L. Feathers, A.M. Ganios, L. Jia, J.M. Miller, and D.A. Thompson. 2015. MERTK signaling in the retinal pigment epithelium regulates the tyrosine phosphorylation of GDP dissociation inhibitor alpha from the GDI/CHM family of RAB GTPase effectors. Exp Eye Res. 140:28–40.

49. Sobota, A., A. Strzelecka-Kiliszek, E. Gladkowska, K. Yoshida, K. Mrozinska, and K. Kwiatkowska. 2005. Binding of IgG-opsonized particles to Fc gamma R is an active stage of phagocytosis that involves receptor clustering and phosphorylation. J Immunol. 175:4450–4457.

50. Sparrow, J.R., D. Hicks, and C.P. Hamel. 2010. The retinal pigment epithelium in health and disease. Curr Mol Med. 10:802–823.

51. Swanson, J.A. 2008. Shaping cups into phagosomes and macropinosomes. Nat Rev Mol Cell Biol. 9:639–649.

52. Thievessen, I., P.M. Thompson, S. Berlemont, K.M. Plevock, S.V. Plotnikov, A. Zemljic-Harpf, R.S. Ross, M.W. Davidson, G. Danuser, S.L. Campbell, and C.M. Waterman. 2013. Vinculin-actin interaction couples actin retrograde flow to focal adhesions, but is dispensable for focal adhesion growth. J Cell Biol. 202:163–177.

53. Tsapara, A., P. Luthert, J. Greenwood, C.S. Hill, K. Matter, and M.S. Balda. 2010. The RhoA activator GEF-H1/Lfc is a transforming growth factor-beta target gene and effector that regulates alpha-smooth muscle actin expression and cell migration. Mol Biol Cell. 21:860–870.

54. Unbekandt, M., D.R. Croft, D. Crighton, M. Mezna, D. McArthur, P. McConnell, A.W. Schuttelkopf, S. Belshaw, A. Pannifer, M. Sime, J. Bower, M. Drysdale, and M.F. Olson. 2014. A novel small-molecule MRCK inhibitor blocks cancer cell invasion. Cell Commun Signal. 12:54.

55. Young, R.W. 1967. The renewal of photoreceptor cell outer segments. J Cell Biol. 33:61–72.

56. Zhao, Z., and E. Manser. 2015. Myotonic dystrophy kinase-related Cdc42-binding kinases (MRCK), the ROCK-like effectors of Cdc42 and Rac1. Small GTPases. 6:81–88.

57. Zihni, C., C. Mills, K. Matter, and M.S. Balda. 2016. Tight junctions: from simple barriers to multifunctional molecular gates. Nat Rev Mol Cell Biol. 17:564–580.

58. Zihni, C., P.M. Munro, A. Elbediwy, N.H. Keep, S.J. Terry, J. Harris, M.S. Balda, and K. Matter. 2014. Dbl3 drives Cdc42 signaling at the apical margin to regulate junction position and apical differentiation. J Cell Biol. 204:111–127.

59. Zihni, C., and S.J. Terry. 2015. RhoGTPase signalling at epithelial tight junctions: Bridging the GAP between polarity and cancer. Int J Biochem Cell Biol. 64:120–125.

60. Zihni, C., E. Vlassaks, S. Terry, J. Carlton, T.K.C. Leung, M. Olson, F. Pichaud, M.S. Balda, and K. Matter. 2017a. An apical MRCK-driven morphogenetic pathway controls epithelial polarity. Nature Cell Biology. 19:1049.

61. Zihni, C., E. Vlassaks, S. Terry, J. Carlton, T.K.C. Leung, M. Olson, F. Pichaud, M.S. Balda, and K. Matter. 2017b. An apical MRCK-driven morphogenetic pathway controls epithelial polarity. Nat Cell Biol. 19:1049–1060.

